# Parallel adaptation and admixture drive the evolution of virulence in the grapevine downy mildew pathogen

**DOI:** 10.1101/2025.05.18.654733

**Authors:** Etienne Dvorak, Thomas Dumartinet, Isabelle D. Mazet, Alexandre Chataigner, Manon Paineau, Dario Cantù, Pere Mestre, Marie Foulongne-Oriol, François Delmotte

## Abstract

- *Plasmopara viticola* is a biotrophic oomycete responsible for grapevine downy mildew, one of the most destructive diseases in viticulture. Breeding for resistant varieties relies on the introgression of partial resistance factors from wild grapes, but virulent strains are rapidly emerging.
- To decipher the genetic bases of the adaptation to plant resistance in *P. viticola*, we carried out a QTL mapping study using two F1 populations segregating for the ability to overcome *Rpv3*.*1, Rpv10* and *Rpv12*. Trajectories of virulence emergence were also compared by conducting a population structure analysis on a panel of diversity.
- We confirmed the position of *AvrRpv3*.*1* and identified the *AvrRpv12* locus, in which strains overcoming *Rpv12* presented large deletions encompassing several RXLR genes. Distinct virulent alleles were selected independently in different winegrowing regions. Unlike this standard case of recessive virulence, partial breakdown of *Rpv10* was determined by a dominant locus, suggesting a suppressor activity. The virulent haplotype exhibits structural rearrangements and an extended effector repertoire. It corresponds to an admixed genomic segment likely originating from a secondary introduction of *P. viticola* into Europe.
- On top of the identification of candidate effectors, these results illustrate the range of evolutionary pathways through which plant pathogen populations can adapt to plant resistances.

## Introduction

Understanding how plant pathogens evolve to overcome host defenses is critical for the effective and sustainable management of crop diseases. Fungal and oomycete pathogens secrete a large number of effectors that promote infection (Koeck et al, 2011; Stassen and Van den Ackerveken, 2011), but some of them act as avirulence (Avr) factors when recognized by resistance (R) proteins (Gassmann and Bhattacharjee, 2012). Effector genes are thus evolving rapidly and evasion of the plant immune response can be achieved via multiple mechanisms, including non-synonymous mutations, deletions and silencing of Avr genes (Petit-Houdenot and Fudal, 2017; Wang et al, 2019). At the population level, the speed of adaptation depends on the ability of pathogens to generate genetic diversity and spread beneficial variants (McDonald and Linde, 2002). In particular, sexual reproduction plays a key role by creating new allele combinations and facilitating gene flow, leading to admixture between populations or even hybridization between species (Ahmed et al, 2012; Leroy et al, 2016; Menardo et al, 2016). Therefore, in addition to the study of genetic determinants of plant-pathogen interactions, maximizing the durability of resistances requires the integration of population genetics approaches to elucidate the factors driving the emergence of virulence.

*Plasmopara viticola* is a biotrophic oomycete causing grapevine downy mildew, one of the most destructive diseases affecting vineyards worldwide. It was introduced from North America to Europe in the 1870s and from there spread to other winegrowing regions around the globe (Fontaine et al, 2021). At the plot level, *P. viticola* populations are large and exhibit a high genotypic diversity (Gobbin et al, 2006). Sexual reproduction occurs each year and outcrossing is ensured by strict heterothallism, resulting in a high heterozygosity rate (Dussert et al, 2019; Maddalena et al, 2020). These elements provide a high evolutionary potential (McDonald and Linde, 2002), as exemplified by the rapid loss of sensitivity to many fungicides (Blum et al, 2010; Delmas et al, 2017).

Due to the high susceptibility of the cultivated Eurasian grapevine *Vitis vinifera*, breeding programs aiming to obtain resistant varieties are based on the introgression of resistances from wild grapes. Most of these genes only provide a partial protection, and their efficiency is variable depending on the environment, the plant physiological state and the genetic background of the variety (Possamai and Wiedemann-Merdinoglu, 2022). Grapevines carrying major R factors trigger a hypersensitive response (HR) upon downy mildew infection, significantly limiting the pathogen’s growth (Casagrande et al, 2011; Wingerter et al, 2021; Juraschek et al, 2022). The best-studied loci encode proteins with nucleotide binding and leucine-rich repeat (NLR) domains, which are known for activating effector-triggered immunity (ETI) (Bellin et al, 2009; Schwander et al, 2012; Venuti et al, 2013). In particular, Rpv1 and Rpv3.1-mediated resistances were proved to be controlled by NLR genes (Feechan et al, 2013; Foria et al, 2020), suggesting they act by recognizing Avr factors produced by *P. viticola*. Most oomycete Avr proteins possess an N-terminal RXLR motif, often associated with a dEER motif. Several RXLR(-like) effectors also share a modular structure mediated by a conserved fold called the (L)WY domain (Boutemy et al, 2011; He et al, 2019). This type of putative effectors is abundant in *P. viticola*, as its genome contains more than 500 RXLR-like genes (Dussert et al, 2019). Oomycete effectors are often encoded in clusters of paralogous genes (Fletcher et al, 2022; Matson et al, 2022). Such genes tend to be located in repeat-rich regions with high rates of duplication and deletion. This genomic compartmentalization is thought to foster an elevated diversity in the effector repertoire, which facilitates the rapid adaptation to plant defenses (Dong et al, 2015; Sánchez-Vallet et al, 2018). Several *P. viticola* secreted proteins have been found to interfere with the plant immune system (Xiang et al, 2016; Combier et al, 2019; Fu et al, 2024), but specific interactions with major grapevine R genes were not investigated until recently.

Over the last few years, studies reported the widespread occurrence of *P. viticola* strains overcoming *Rpv3*.*1*, and the recent breakdown of *Rpv10* and *Rpv12*, two loci introgressed from the East Asian species *Vitis amurensis* (Heyman et al, 2021; Wingerter et al, 2021; Paineau et al, 2022). Thanks to the last advancements on the pathogen’s genomics, the first *P. viticola* Avr locus was identified in a genomewide association study (GWAS) (Paineau et al, 2024). Virulence towards *Rpv3*.*1* is associated with the absence of a tandem of secreted DEER proteins, and the locus presents an important allelic diversity. The interaction between *P. viticola* and *Rpv3*.*1*-carrying plants thus fits a gene-for-gene relationship, in which resistance is mediated by the recognition of specific pathogen effector(s) (Rouxel and Balesdent, 2010).

This raises the question of whether the newly observed virulences in *P. viticola* also result from the loss of effector genes, allowing escape from host immunity. Understanding the evolutionary trajectories of pathogen populations is also crucial to inform the deployment and management of new resistant grapevine varieties. It remains to be determined if virulences arise from the selection and subsequent spread of a single haplotype carrying a favorable variant, or rather from independent mutations at the same loci in different subpopulations.

In this study, we aimed to characterize in parallel the genetic bases of the adaptation to two major grapevine resistances in *P. viticola*. Using quantitative trait locus (QTL) mapping in two F1 populations, we identify the genomic regions linked to virulence towards *Rpv12* and *Rpv10*. We show that both loci are dynamic effector-rich regions, yet their modes of inheritance differ, the first being recessive and the second dominant. The structure of *P. viticola* populations suggests that adaptation to *Rpv12* has occurred independently across multiple winegrowing regions, while virulence towards *Rpv10* was likely acquired through a single admixture event. These results highlight the diverse evolutionary pathways through which a specialized pathogen can adapt to plant resistances.

## Material and methods

### Pathogen strains

The generation of the two F1 mapping populations was described in Dvorak et al (2025). The first cross involved strains Pv412 11 and Pv2543 1 (N=162) with contrasting pathotypes on Rpv3.1 and Rpv12 hosts (Table 1) (Paineau et al, 2022). The second progeny was the result of a cross between Pv412 11 and Pv1419 1 (N=189), the latter being able to overcome *Rpv10* (Table 1). For brevity, these two F1 populations are hereafter referred to as 412×2543 and 412×1419.

**Table 1.** Origin and pathotype of parent strains.

|  | Origin | Rpv3.1 | Pathotype on Rpv10 | Rpv12 |
| --- | --- | --- | --- | --- |
| **Pv412_11** | Canton of Ticino, Switzerland | virulent | avirulent | avirulent |
| **Pv2543_1** | Baranya county, Hungary | avirulent | avirulent | virulent |
| **Pv1419_1** | State of Baden-Württemberg, Germany | virulent | virulent | avirulent |

In addition, strain cPv44 1 from the 412×1419 population was backcrossed to the *Rpv10*-breaking parent Pv1419 1 in order to test the dominance of this trait. Crossing, maturation and oospore retrieval were performed as described in Dvorak et al (2025). The progeny is referred to as the BC-1419 population (N=51).

### Phenotyping experiments

#### Plant material

*P. viticola* strains were phenotyped on a set of grapevine cultivars carrying different Rpv factors. We used cv. ‘Cabernet-Sauvignon’ (susceptible reference), cv. ‘Regent’ (*Rpv3*.*1*), cv. ‘Muscaris’ (*Rpv10*) and cv. ‘Fleurtai’ (*Rpv12*). The BC-1419 population was also phenotyped on an additional cultivar called ‘Solaris’ that carries *Rpv10*.

#### Plant inoculation

F1 progenies were inoculated on leaf discs of susceptible and resistant plants in separate experiments. Due to the large number of strains to phenotype, progenies were divided in half and inoculated in two parts, ten days apart, with a set of reference strains repeated in all experiments. Grapevine scions grafted onto the *Vitis berlandieri x riparia* ‘SO4’ rootstock were grown in a greenhouse without chemical treatment and under natural photoperiod conditions for 6 to 7 weeks. Leaf discs preparation, inoculation and incubation were performed as described previously (Paineau et al, 2022). Briefly, strains were initially propagated on detached leaves of cv. ‘Cabernet-Sauvignon’. One day before the experiment, infected leaves were gently rinsed with distilled water to ensure the production of fresh sporangia for the inoculation the next day. Sporangia from each strain were suspended in sterile water and concentrations were adjusted to 10^5^ sporangia per ml using a Scepter 2.0 portable particle counter (Millipore). Leaf discs were excised from the fourth leaf below the apex, and placed on wet filter paper in a 12×12 cm Petri dish. Each suspension was sprayed on one Petri dish containing 5 discs per variety. For a given interaction, each disc came from a different plant. Plates were sealed with plastic film, and then incubated for 6 days in a growth chamber at 18°C with a 12:12 photoperiod. The BC-1419 population was phenotyped with the same method by inoculating 8 discs of ‘Cabernet-Sauvignon’, ‘Muscaris’ and ‘Solaris’, and 4 discs of ‘Fleurtai’.

#### Traits measurement

The percentage of sporulation area was calculated on high-resolution pictures taken at 6 days post-inoculation (dpi) using an in-house image analysis program (code available on https://gitlab.com/grapevinedownymildew/notebookimageanalysis). On the same pictures, necrotic lesions were visually assessed using an ordinal scale based on their size and appearance (Fig.S1). Discs received a score between 1 (large non-specific necrotic speckles) and 5 (small dark lesions indicative of an efficient HR). Complete absence of necrosis was scored as 0.

#### Statistical analyses

Analyses were conducted using R v4.1.3. Broad-sense heritabilities (H2) were calculated for each trait on the different inoculated hosts using the function H2cal implemented in r/inti v0.6.6 (Lozano-Isla, 2024). As F1 offspring of the same population were phenotyped in two parts, we fitted a linear mixed model with the experiment (Exp) treated as a random intercept effect, and the percentage of sporulation area (Spo) for each individual (Ind) were calculated as best linear unbiased estimations using r/emmeans v1.10.2 (Lenth, 2024). This was done separately for each inoculated variety. The model was noted as *Spo*_*ijk*_ = *µ* + *Ind*_*i*_ + *u*(*Exp*)_*j*_ + *ϵ*_*ij*_ with 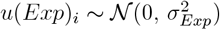 and *ϵ*_*ijk*_ ∼ 𝒩 (0, *σ*^2^).

For the study of the BC-1419 population, we tested the effect of the genotype (Geno), the inoculated host (Host), and their interaction on the sporulation area (Spo) by fitting a linear mixed model with the Ind variable treated as a random intercept effect. The model was noted as *Spo*_*ijkl*_ = *µ* + *Geno*_*i*_ + *Host*_*j*_ + (*Geno* : *Host*)_*ij*_ + *u*(*Ind*)_*k*_ + *ϵ*_*ijkl*_ with 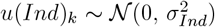 and *ϵ*_*ijkl*_ ∼ 𝒩 (0, *σ*^2^). Pairwise comparisons between groups were performed with r/emmeans.

#### QTL mapping

Parental linkage maps were built based on targeted genotyping-by-sequencing of SNP markers and were previously presented in Dvorak et al (2025). Triploid strains were not included in the following analyses. Thus, 6 individuals were removed in the 412×1419 progeny and 7 in 412×2543, resulting in respectively 183 and 155 genotypes effectively used.

Single interval QTL mapping was performed using r/qtl v1.60 (Arends et al, 2010). A square-root transformation was applied to measures of sporulation area to normalize their distribution. LOD scores were calculated using a normal model and Haley-Knott regression. Significance levels were computed with 1000 genome scan permutations. QTL boundaries were determined by calculating credibility intervals with the bayesint function. The percentage of variance explained by each QTL was calculated using the fitqtl command. Composite interval mapping was also tested using the same package but did not reveal additional QTLs and did not change credibility intervals.

### Analysis of putative effector genes in the QTLs

All analyses were conducted using the most recent assembly of reference strain Pv221 1 (Paineau et al, 2024). Genes located in QTL intervals were checked for the presence of signal peptides (SP), RXLR and/or dEER motifs, as well as LWY domains, a structural fold associated with oomycete effectors. Secreted proteins were predicted using SignalP 5.0 (Almagro Armenteros et al, 2019). LWY domains were searched with HMMER 3.2 (hmmer.org) as described in Dussert et al (2019), using the HMM profile from Boutemy et al (2011).

Structural homology searches were conducted with Phyre2 (Kelley et al, 2015). Predictions of protein structure were performed using AlphaFold2 (Jumper et al, 2021) as implemented in ColabFold v1.5.2 (Mirdita et al, 2022) using default settings. Visualization and superimposition of protein structures were visualized and in UCSF Chimera X 1.5 (Goddard et al, 2018). The experimental structure of the *Phytophthora sojae* PSR2 effector was retrieved from the Protein Data Bank (https://www.ebi.ac.uk/pdbe/pdbe-kb/proteins/E0W4V5).

Expression of the genes present in the identified QTLs was checked during plant infection by the reference avirulent strain Pv221 1. Transcript analysis was conducted using processed RNA-sequencing data available from Dussert et al (2019) (SRA BioProject PRJNA329579).

### Haplotype-resolved assembly of parent strain Pv1419 1

We benefited from a *de novo* assembly of the parent strain Pv1419 1. High-molecular weight DNA was extracted from sporangial tissues using the same protocol described in Dussert et al (2019). DNA fragments were sequenced on a PacBio Sequel II system, yielding long high-fidelity reads. They were assembled into diploid pseudochromosomes using the haplotype-aware pipeline HaploSync (Minio et al, 2022). Assembly procedure, quality assessment and annotation are detailed in Methods S1.

### Genotyping of the backcross population by amplicon length polymorphism

After the detection of a QTL in the 412×1419 F1 progeny, we aimed to follow its inheritance in the BC-1419 population. Indels were identified between the haplotypes carried by Pv1419 1 at the QTL: one at each edge of the physical interval, and one co-segregating with the QTL peak. For each genotype, different profiles of amplicon lengths were generated by PCR and visualized by electrophoresis on agarose gels (Methods S2). Primer sequences and annealing temperatures are indicated in Table S2.

### Whole genome sequencing of additional strains

We sequenced 41 new *P. viticola* strains, including three North American isolates belonging to clade *aestivalis*, the only species present worldwide (Fontaine et al, 2021). Additionally, whole genome sequencing (WGS) data were obtained for six F1 individuals and one BC-1419 strain, which enabled the phasing inherited variants in the QTLs we detected. DNA was extracted from sporangial tissues following a CTAB protocol adapted from Möller et al (1992) and previously detailed in Dvorak et al (2025). Libraries were prepared using an Illumina DNA TruSeq kit. Sequencing was performed at the GeT-PlaGe facility (Toulouse, France) with a NovaSeq6000 to produce 2×150 bp paired-end reads. This applied to all samples except Canadian isolates, for which DNA was sequenced by Beckman Coulter Genomics (Grenoble, France) on an Illumina HiSeq 2000 sequencer (2×100 bp paired-end reads).

### Population structure analysis

We constructed a panel of sequences that included strains of various pathotypes collected from different European regions as well as other continents (Data S1). All samples that have a number as a suffix (‘-1’ or ‘-11’) were derived from monosporangium isolation as described in Paineau et al (2022). Short DNA reads were obtained either in previous studies or for the present one (Data S1). In total, 56 wild strains and 7 F1 or BC-1419 individuals were included.

The population structure was investigated using SNP data from wild strains. Read mapping, variant calling and filtration are detailed in Methods S3. Variants were pruned to retain those in approximate linkage equilibrium with PLINK 1.9 (Chang et al, 2015). The parameters used were a window size of 50 variants with a shift of 10 variants at each step, and a r2 threshold of 0.1. After pruning, 228,318 SNPs remained. Using PLINK, KING-robust kinship coefficients were computed to check for closely related samples. A Principal Component Analysis (PCA) was performed using r/adegenet v2.1.10 (Jombart and Collins, 2015). Population clusters and individual ancestries were then inferred using ADMIXTURE v1.3 with unsupervised analysis (Alexander and Lange, 2011). The program was run for K=1 to K=5 with 200 bootstrap replicates each time, and cross-validation (CV) errors were calculated for each K.

Ancestry segments in admixed individuals were identified using MOSAIC v1.5.1 (Salter-Townshend and Myers, 2019). As this approach required variant phasing, we used only a subset of SNPs that were fixed in each “ancestral” population (homozygous reference allele in the pure European subset, homozygous alternate allele in the North American population). Genotypes of admixed strains were then phased using Beagle v5.2 (Browning et al, 2021) with the two mentioned sub-populations set as references. Standard parameters were used, except for sliding windows that were set to 1 Mb with 0.1 Mb overlaps. Local ancestry along the genome was finally estimated using MOSAIC with default parameters and rephasing enabled.

## Results

### Mapping of three major QTLs involved in resistance breakdown

Two biparental populations were derived from the cross of *P. viticola* strains exhibiting contrasting pathotypes on three grapevine resistance factors (Table 1). Phenotyping on resistant cultivars revealed segregation on *Rpv3*.*1* and *Rpv12* plants in the 412×2543 progeny, and segregation on *Rpv10* plants in the 412×1419 progeny. One major QTL was detected for each R gene, in different parental linkage maps (Fig. S2).

We first confirmed the position of the *AvrRpv3*.*1* locus, which was previously identified by a genomewide association study (GWAS) (Paineau et al, 2024). In the 412×2543 population, 88 out of 162 individuals did not trigger HR (average necrosis score = 0), suggesting that virulence towards *Rpv3*.*1* segregated with a 1:1 ratio (Chi-squared test, p = 0.31) (Fig. S3A). A single major QTL was detected at the edge of LG11 in the linkage map of the avirulent parent Pv2543 1 (Fig. S3B-C), with a credibility interval of 5.1 cM. The QTL explained 46.1% of the variation in sporulation area and 87.1% of the variation in necrosis score.

The genetic interval corresponded to a 148-kb long physical segment encompassing a tandem of effector genes, Primary 000014F.g164 and Primary 000014F.g165 (P14g164 and P14g165). Read depth analysis showed that P14g164 and P14g165 were partially or totally deleted in both haplotypes of the virulent parent Pv412 11 (Fig. S3D). These genes were present only in the avirulent haplotype of Pv2543 1, thus confirming their role as Avr factors recognized by Rpv3.1 (Paineau et al, 2024).

### Virulence towards Rpv12 is associated with the loss of RXLR genes

The 412×2543 F1 population also segregated regarding the interaction with *Rpv12*. The sporulation area varied along a continuum from 0 to 11% (Fig. 1A). Trait heritabilities were lower than those observed in the interaction with *Rpv3*.*1*, with H2=0.40 for the sporulation area and 0.35 for the necrosis score. High levels of sporulation tended to be associated with light necrosis surrounding sporulation spots (average necrosis score < 3). This did not correspond exactly to the parental phenotype, as Pv2543 1 infected *Rpv12* plants without triggering HR.

**Fig. 1:**
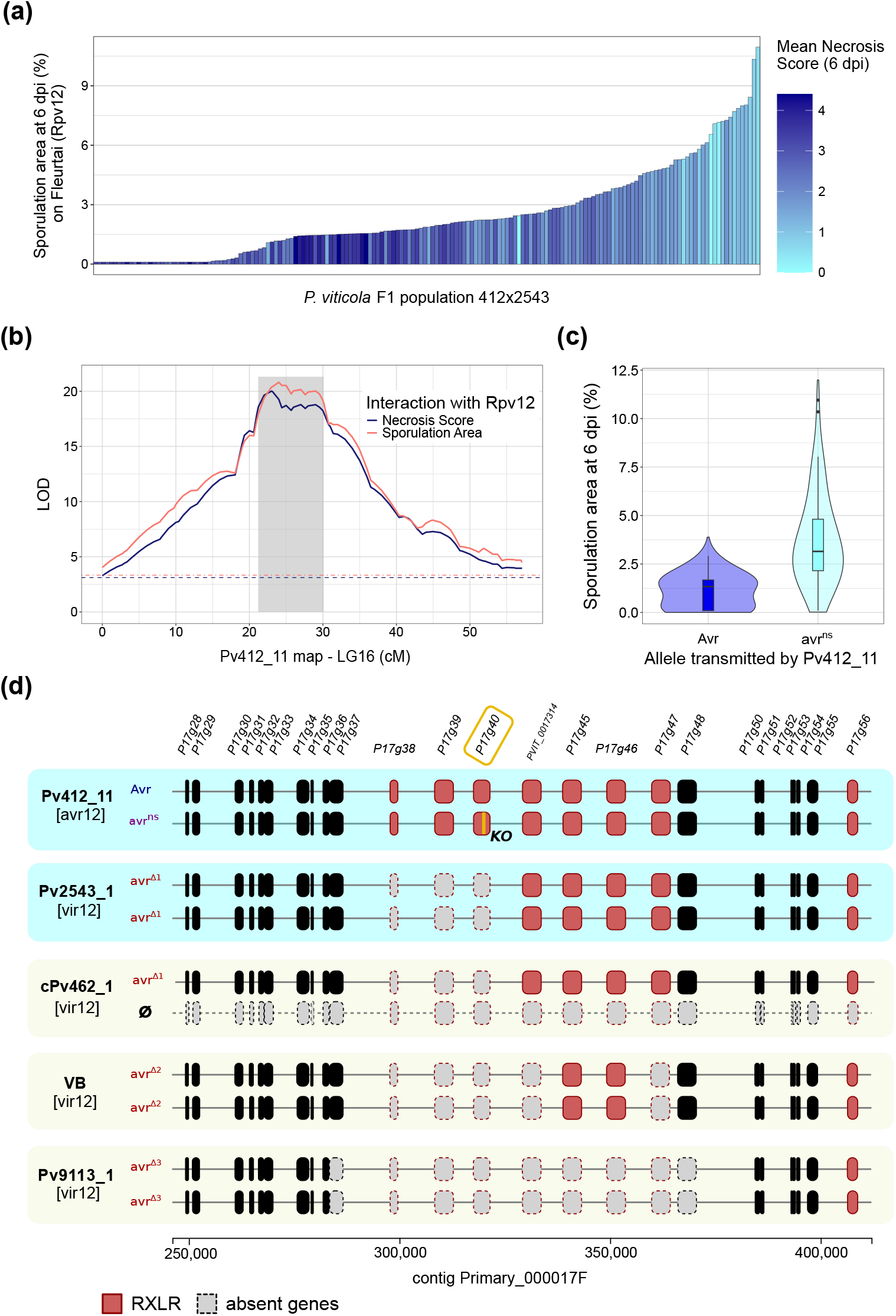
Mapping and characterization of the *AvrRpv12* locus in a biparental population of *P. viticola*. **(a)** Phenotypes distribution in the 412×2543 F1 progeny (N=162) on cv. ‘Fleurtai’ (*Rpv12*). **(b)** QTL mapping of *Rpv12*-breakdown in the linkage map of the avirulent parent Pv412 11. The gray area indicates the credibility interval of the QTL. Dashed lines indicate the LOD significance thresholds determined using 1000 permutations. Results on other linkage groups and other cultivars are available in Fig. S2. **(c)** Distribution of the sporulation area on *Rpv12* depending on the marker allele at the QTL peak. Horizontal lines in the boxplots signal the 25^th^, 50^th^ and 75^th^ percentiles. **(d)** Allelic configurations of parent strains (blue boxes) and other virulent strains (beige boxes) in the physical interval of the QTL. For each strain, the two haplotypes are represented. Genes are colored in black or red if they possess a RXLR motif. Dashed gray boxes represent absent genes. Deletion patterns were deduced from read depth (Fig.S5). In the Pv412 11 haplotype associated with virulence (avr^NS^), an orange stroke signals the 1-nt indel causing a premature stop codon in P17g40. Strain cPv462 1 is an aneuploid offspring that lacks the copy of chromosome 16 transmitted by Pv412 11.

A single major QTL was detected in LG16 in the linkage map of the avirulent parent Pv412 11, with a similar profile for the two measured traits and a genetic interval of 8.9 cM (Fig. 1B). No QTL was detected on *Rpv12* plants in the linkage map of the virulent parent (Fig. S2). The variance explained by the Pv412 11 QTL was 28.9% for the sporulation area and 31.7% for the necrosis score. The distributions of sporulation area between the two alleles partly overlapped (Fig. 1C). As Pv412 11 itself showed a quasi-absence of sporulation, these two alleles were seemingly revealed by the cross with the virulent strain.

We analyzed the physical region underlying the QTL which corresponded to a 157-kb long segment in the reference genome (Fig. 1D). This interval comprises 25 coding sequences. One of them (PVIT 0017314) was incorrectly annotated so we used the name given in the first version of the *P. viticola* genome annotation instead (Dussert et al, 2019). Most genes have no functional annotation (Data S2). However, eight of them encode proteins that contain a secretion peptide (amino acids 1-20) and a RXLR motif (aa 49-52). Thus, we focused primarily on these genes as they are likely to encode cytoplasmic effectors. Transcriptome analysis showed that they were all expressed during plant infection by the avirulent reference strain Pv221 1. Their protein sequences vary from 608 to 1481 aa. All these RLXR genes are highly similar (>50% of amino acid identity) and belong to a family of 18 genes located in a 300-kb segment around the QTL (Fig. S4). Phylogenetic analysis showed that related sequences tend to be physically close (Fig. S4). Two genes, PVIT 0017314 and P17g45, have an increased read depth in both parents which is probably due to additional copies compared to the reference genome (Fig. S5). These elements suggest that this region is subjected to repeated tandem duplications generating copy number variation (CNV) and paralogous sequences.

Read depth analysis in the virulent parent strain Pv2543 1 revealed a large homozygous deletion containing three RXLR genes: P17g38, P17g39 and P17g40 (Fig. 1D and Fig. S5). They code for proteins sharing structural homology with known oomycete effectors, such as PSR2 from *P. sojae* and RXLR12 from *Phytophthora capsici* (best hits obtained with Phyre2). HMM search also highlighted the presence of repeated LWY folds in the three proteins (respectively 3, 10 and 9 modules). We modeled the structure of P17g40 using AlphaFold2 and divided it based on the identified LWY modules, which revealed a clear and complete overlap with the PsPSR2 modules (Fig. S7).

Due to their absence in Pv2543 1, the 412×2543 offspring inherited only one copy of these genes. The other parent, Pv412 11, presented an important heterozygosity in their coding sequences, with respectively 9, 45 and 13 non-synonymous mutations. Whole-genome sequencing of three 412×2543 offspring enabled us to phase variants in these genes. P17g38 and P17g39 share respectively 98.5% and 98.4% nucleotide sequence identity between the two alleles, while P17g40 sequences are 99.2% identical. However, for this last gene, a 1-nt deletion is present in the allele associated with higher sporulation. This frameshift mutation leads to a truncated protein at the position 795/1298 (Fig. 1D). We noted the haplotype avr^ns^ because of this nonsense mutation in a putative effector.

Given the major variations affecting RXLR genes in this locus, we hypothesized than one or several of them could correspond to an Avr gene recognized by *Rpv12*. We checked their status in other *P. viticola* strains. We found that all strains virulent towards *Rpv12* were affected by large deletions in the locus, but their lengths varied depending on the location of origin (Fig. 1D and Fig. S5). Hungarian strains presented the same deletion as Pv2543 1 (allele noted avr^Δ1^). In the Swiss VB strain described in Wingerter et al (2021), 5 RXLR genes were absent (avr^Δ2^). Finally, virulent Italian strains (like Pv9113 1) lacked 7 out of the 8 RXLR genes in the QTL (avr^Δ3^). We also identified one Hungarian strain (Pv2963) that was fully avirulent and appeared hemizygous at the *AvrRpv12* locus (Fig. S5). This observation is consistent with a recessive virulent allele. Besides, all *Rpv12*-breaking strains presented long runs of homozygosity (ROH) around the deletions (Fig. S6). These ROHs extended from 400 to around 700 kb and could be the result of a recent selective sweep.

One genotype in the 412×2543 F1 progeny provided additional clues that the QTL corresponded to *AvrRpv12*. Strain cPv462 1 is aneuploid (Dvorak et al, 2025) and lacks the copy of chromosome 16 normally transmitted by the avirulent parent Pv412 11 (Fig. 1D). This strain did not trigger any necrosis on *Rpv12* plants (average necrosis score = 0). The absence of HR was further verified by inoculating additional ‘Fleurtai’ leaf discs by droplet and observing the discs over one week. This confirmed that avirulence towards *Rpv12* is entirely carried by chromosome 16. Together, these findings led us to consider RXLR genes in the QTL as strong AvrRpv12 candidates.

### A major QTL determines the partial breakdown of *Rpv10*

The 412×1419 F1 population (N=189) was used for QTL detection in the interaction with *Rpv10*. A large part of the offspring showed little to no sporulation on cv. ‘Muscaris’ (Fig. 2A). The median sporulation area was only 0.5%, with the most aggressive strains reaching 6% of sporulation area. For comparison, the median sporulation area reached 7.2% on the susceptible cultivar ‘Cabernet-Sauvignon’. Necrotic lesions were present on almost every leaf discs (Fig. 2A). Sporulation area and necrosis score showed a heritability of 0.55 and 0.59 respectively.

**Fig. 2:**
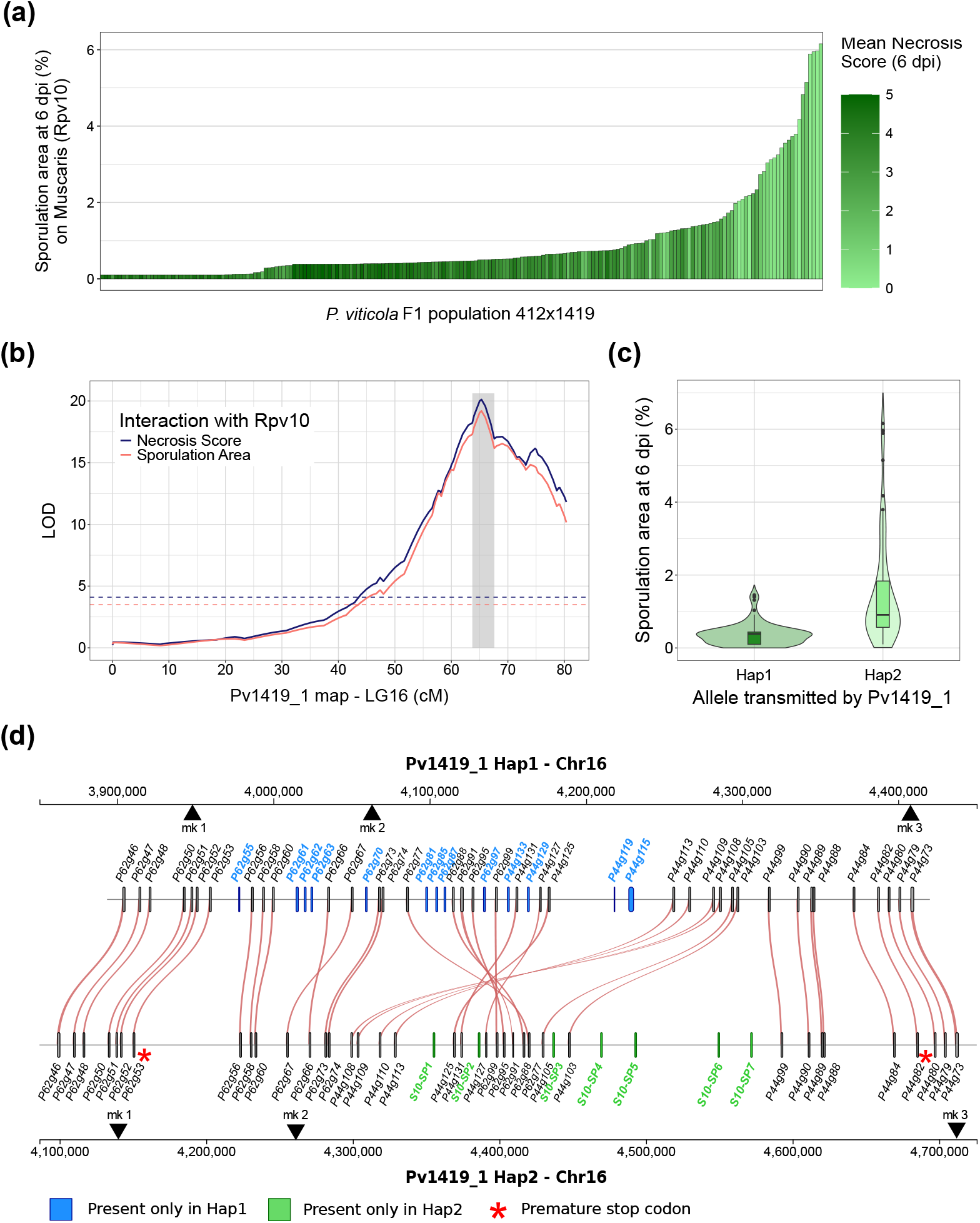
Identification of a locus determining the partial breakdown of Rpv10 in a biparental population of *P. viticola*. **(a)** Phenotypes distribution in the 412×1419 F1 progeny (N=189) on cv. ‘Muscaris’ (*Rpv10*). **(b)** QTL mapping of *Rpv10*- breakdown in the linkage map of the virulent parent Pv1419 1. The gray area indicates the credibility interval of the QTL. Dashed lines indicate the LOD significance thresholds determined using 1000 permutations. Results on other linkage groups and other cultivars are available in Fig. S2. **(c)** Distribution of the sporulation area on *Rpv10* depending on the marker allele at the QTL peak. Horizontal lines in the boxplots signal the 25^th^, 50^th^ and 75^th^ percentiles. **(d)** Comparison of the two haplotypes of Pv1419 1. The avirulent haplotype is represented at the top (Hap1). The haplotype associated with partial breakdown of *Rpv10* is at the bottom (Hap2). Genes encoding putative secreted proteins are represented and linked in red when they are present in both haplotypes. Sequences present only in one haplotype are colored in blue (Hap1) or green (Hap2). Genes are named after their annotation in the reference assembly of Pv221 1. Coding sequences of Hap2 that are absent from the reference genome were named S10-Secreted Protein (SP) 1 to 7. Red asterisks signal genes containing premature stop codons. Black triangles indicate the positions amplified by PCR for the genotyping of the BC-1419 backcross progeny (Table S2).

A single major QTL was detected on LG16 in the linkage map of the virulent parent Pv1419 1 (Fig. 2B). The QTL could be narrowed down to a genetic interval of 2.4 cM. It explained 27.5% of the variance in sporulation area and 35.4% of the variance in necrosis score. No QTL was found on other grapevine cultivars or in the other parental map (Fig. S2). In our setting, a QTL could be detected if the parent strain were heterozygous, which suggested that the virulence of Pv1419 1 was due at least in part to a dominant or co-dominant locus. Because the HR was still activated in cv. ‘Muscaris’, we refer to the observed phenotype as a partial breakdown of *Rpv10*.

We then explored the physical segment underlying the identified QTL. It was found on the same linkage group as the *AvrRpv12* locus but the two regions are clearly distinct: the two intervals are located on both sides of the centromere (Dvorak et al, 2025) and separated by at least 1 Mb. Despite the limited genetic interval, the physical segment was large: at least 537 kb, spread over two contigs (Primary 000044F and Primary 000062F). This was mainly due to the absence of recombination in a 248-kb long region which coincided with the peak of the QTL. A large fraction of the locus length (24%) was annotated as repeats, mostly corresponding to Copia-like long terminal repeat TEs.

A striking feature of the QTL interval was a strong enrichment in genes coding for secreted proteins. It contains 120 genes, 50 of which encode an N-terminal SP. This represents 42% of the coding sequences in the interval, compared to 9.7% in the entire genome. Given the large size of the region, we focused primarily on these genes because of their potential role in the interaction with the host. All genes except one were expressed during plant infection by the reference avirulent strain Pv221 (Data S3). Almost all of them code for proteins of similar lengths (300-400 aa) with typical oomycete effector features. They lack the RXLR motif, but possess an N-terminal DEER sequence (typically in aa positions 55-58) and exhibit repeated LWY domains (1 to 4, mostly 3, Data S3). Therefore, the QTL constitutes a hotspot of putative effectors that could be involved in the partial breakdown of *Rpv10*. However, we were still limited in our comprehension of the genomic region due to the lack of an assembly of the distinct haplotypes.

### Extensive structural variation between the two parental haplotypes

To investigate differences between haplotypes, HiFi reads of strain Pv1419-1 were used to build a chromosome-scale assembly (2n=34) in which both haplotypes were fully represented. We used the SNPs included in the linkage map to associate haplotypes with their corresponding phenotypes and verify that the phasing was correct along the QTL.

The two parental haplotypes carried by Pv1419 1 presented extensive structural variation (Fig. 2D). The first haplotype (Hap1) corresponded to the fully avirulent phenotype. It stretched over 507 kb, and its gene content and order were identical to the reference strain Pv221. By contrast, the second haplotype (Hap2) was highly divergent and considerably larger (617 kb). The central part of the segment presented a large inversion that affected around 213 kb in Hap1 and 122 kb in Hap2. The position of this structural variation coincided with the non-recombining region in the linkage map. This large-scale inversion probably explains why crossovers were prevented in such a long interval. The gene content also differed: 13 secreted protein genes are missing in Hap2 and were not found anywhere else in the chromosome (Fig. 2D). Interestingly, this is counter-balanced by the presence of 7 new genes that are exclusive to Hap2 (genes labeled SP1 to SP7), four of which are located in a 100-kb insertion located near the inversion. All seven genes code for secreted proteins that possess an N-terminal DEER motif and 3 repeated LWY domains (Data S4). They correspond to additional copies or paralogous sequences of neighboring genes (between 59 and 94% of aa identity) (Data S4).

We also examined punctual variants affecting genes conserved in both haplotypes. Two genes, P44g82 and P62g53, were affected by premature stop codons in Hap2 (red asterisks in Fig. 2D). Between haplotypes, protein sequences were 97.3% identical on average (min. 93.5%, max. 100%) (Data S4). Altogether, the haplotype responsible for the partial breakdown carried important punctual and structural variations.

### Partial virulence towards *Rpv10* is dominant in a backcross population

By contrast with other QTLs, the one involved in *Rpv10* breakdown was not recessive. Thus, we provisionally designate the locus Suppressor of Avirulence towards Rpv10 (*S-AvrRpv10*). For brevity, we labeled the allele associated with full avirulence “s10” and the active allele “S10”. The 412×1419 F1 population was only composed of S10/s10 and s10/s10 individuals. Thus, the characterization of the homozygous S10/S10 genotype on *Rpv10* was crucial to determine if the S10 allele was fully dominant or co-dominant.

We generated and phenotyped the backcross population BC-1419 (N=51). Strains were genotyped by PCR-based amplicon length polymorphism (Data S5). Three markers were designed, with one cosegregating with the QTL peak (position “mk2” on Fig. 2D). The observed segregation was in accordance with the expected 1:2:1 ratio (18:26:12, p=0.22).

The genotype at the *S-AvrRpv10* locus had no effect on the susceptible cultivar ‘Cabernet-Sauvignon’ (Fig. 3). Conversely, all strains were fully avirulent towards *Rpv12*. On *Rpv10* cultivars ‘Muscaris’ and ‘Solaris’, the sporulation area was higher in strains carrying one or two copies of the S10 allele. However, there was no difference of aggressiveness between heterozygous S10/s10 and homozygous S10/S10 strains (Fig. 3). Weak necrosis was still observed on discs inoculated with S10/S10 individuals (average necrosis score = 2.30 on ‘Muscaris’ and 2.15 on ‘Solaris’). Therefore, we conclude that the S10 allele is fully dominant and that its effect is specific to the interaction with *Rpv10*. It is however not sufficient to completely prevent the immune response in *Rpv10* cultivars, as indicated by the presence of necrosis for all genotypes.

**Fig. 3:**
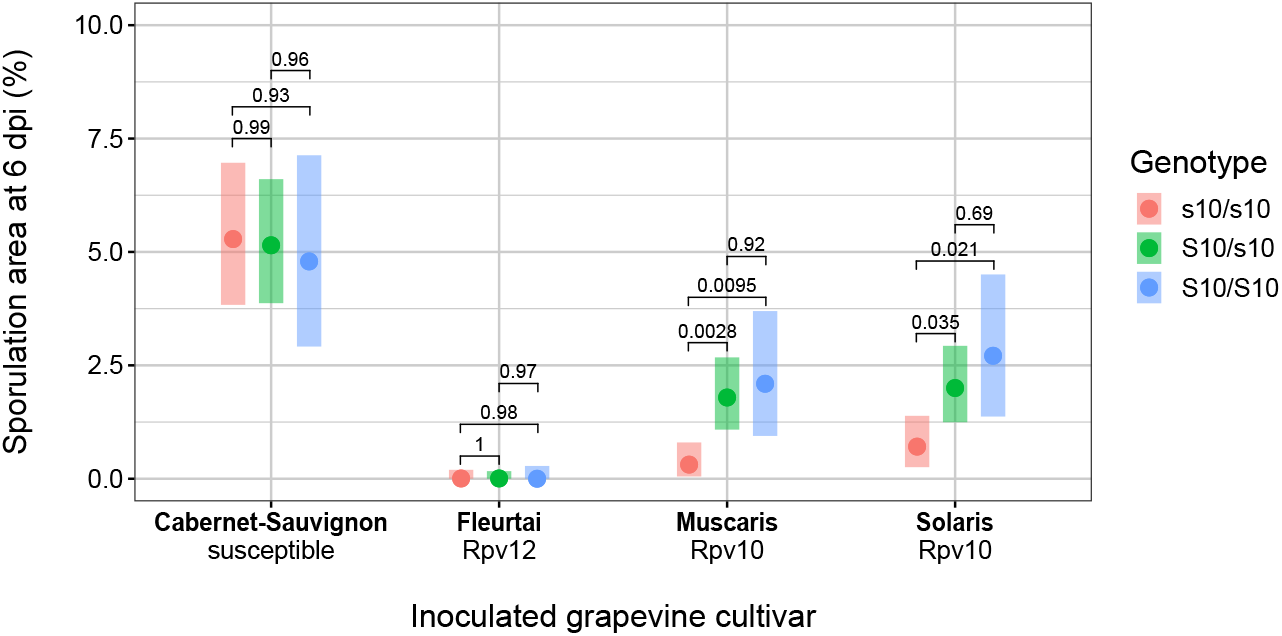
Phenotyping of a *P. viticola* backcross population segregating for partial virulence towards *Rpv10*. The BC-1419 population (N=51) was obtained by crossing S10/s10 strains, thus generating three genotypes at the QTL in the progeny. Offspring were genotyped using three markers by PCR-based amplicon length polymorphism (Table S2). The genotypes shown were obtained using the marker co-segregating with the peak of the QTL in the F1 population (Fig. 2D). Dots indicate the adjusted means and colored bars represent confidence intervals (*α*=0.05). Tukey’s adjusted p-values were calculated for pairwise comparisons with r/emmeans (Lenth, 2024). Full genotyping results are available in Data S5.

### Population structure points to different scenarios of adaptation to *Rpv* genes

We took advantage of WGS data obtained from a panel of strains of various origins and pathotypes (Data S1) to explore the population structure of *P. viticola* in light of our results above. In particular, we aimed to elucidate if the adaptation to each resistance factor in Europe stemmed from a common genetic background.

First, we performed a PCA based on a set of 228,312 SNPs (Fig. 4A). The first PC separated North American strains from the rest of the world while the second PC mostly corresponded to an east-west gradient in Europe. Genetic clustering with the ADMIXTURE program was in accordance with the geographical structuration suggested by PCA (Fig. 4B). The CV error was lowest for K=3. American strains clustered together consistently (cluster 1). With K=3, French and Spanish strains tended to be included in cluster 2 while cluster 3 regrouped most Italian and Hungarian strains.

**Fig. 4:**
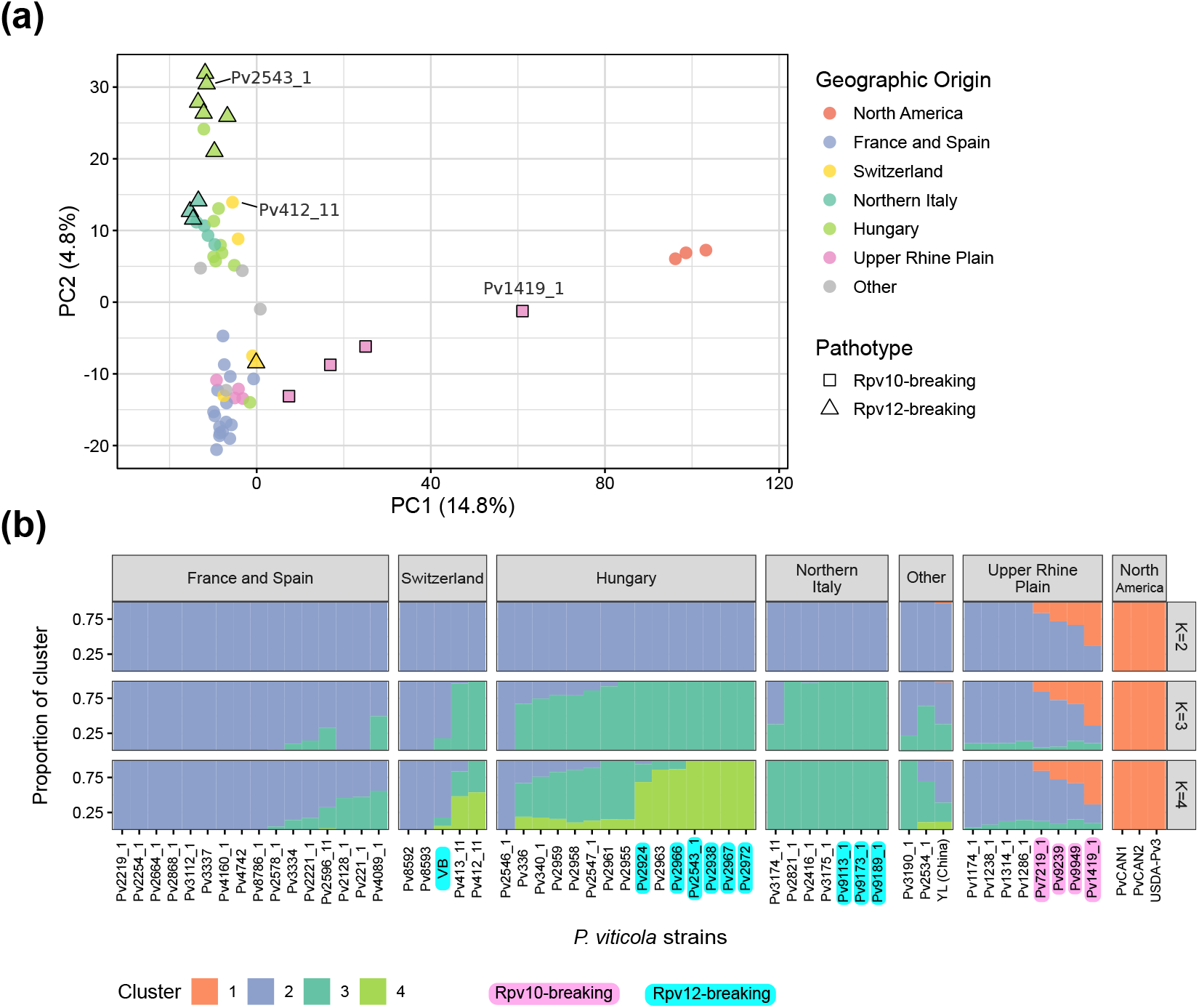
Population structure of *P. viticola* strains with various origins and pathotypes. Analyses were based on 228,318 SNPs in a panel of 57 strains. The Upper Rhine Plain is a winegrowing region at the French-German border. **(a)** Principal Component Analysis. The first two principal components (PC) are represented, with the percentage of variance explained indicated in parentheses. Strains are colored according to their geographical origin, and shapes indicate pathotypes of interest. The positions of parent strains used for QTL mapping are signaled. **(b)** Unsupervised genetic clustering performed with the ADMIXTURE program. Bar plots represent estimated genome ancestry fractions for each strain. Results are shown for 2 to 4 clusters (K). Cross-validation error was lowest with K=3. Strains are separated by geographical origin. The “Other” category regroups non-European strains collected (from left to right) in Georgia, Lebanon and China. Strains overcoming *Rpv10* or *Rpv12* are highlighted in pink and cyan, respectively. Plot made with r/starmie v0.1.3.

**Fig. 5:**
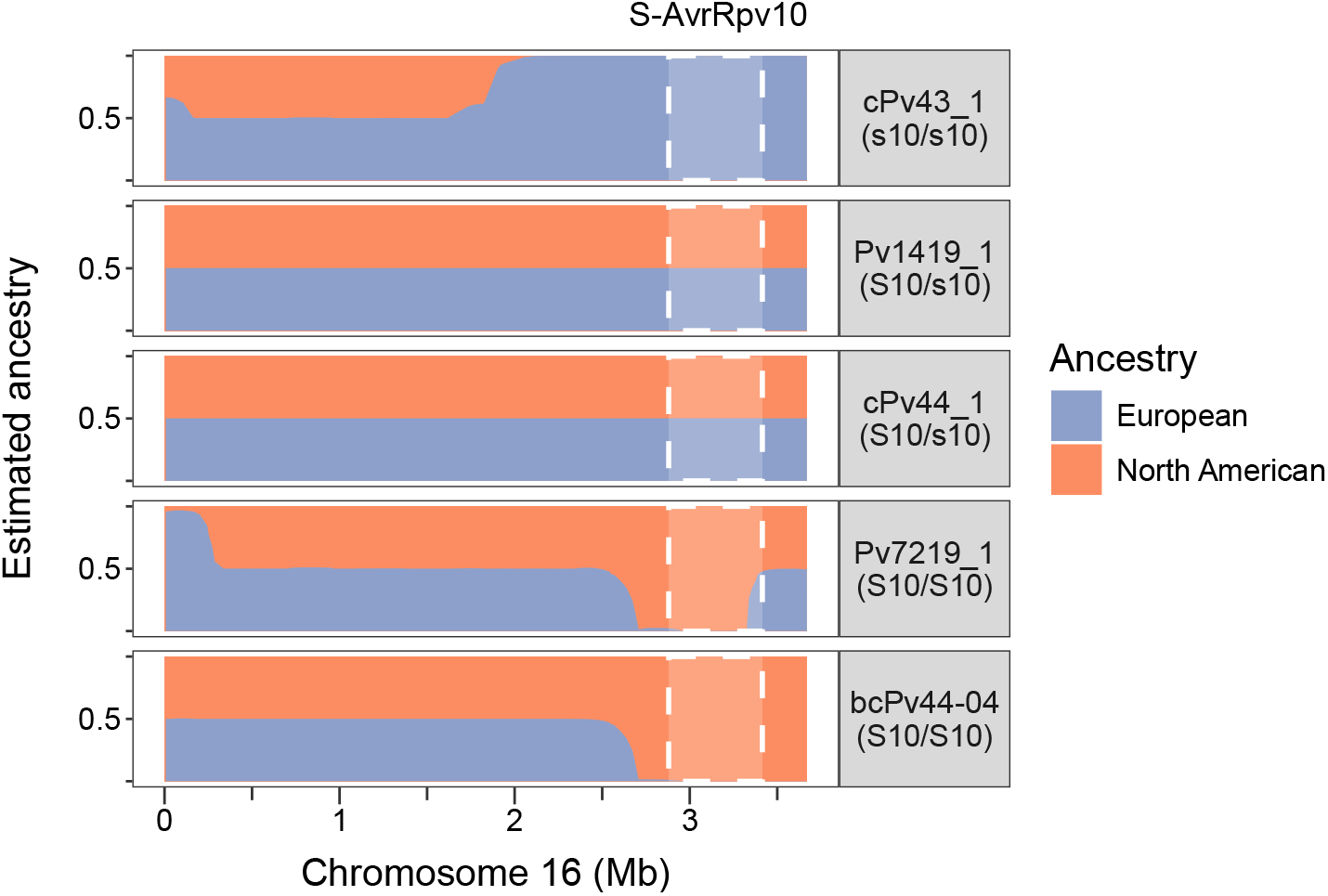
Ancestry segments in admixed *P. viticola* strains. Estimation of local ancestry was conducted on chromosome 16 using a subset of 3876 SNPs. Non-admixed European and North American samples were set as ancestral populations. Proportions of ancestry were estimated for natural admixed strains as well as descendants of Pv1419 1 (F1: cPv43 1 and cPv44 1, backcross: bcPv44-04). The *S-AvrRpv10* QTL limits (Fig. 2) are indicated by white dashed lines. Five strains of different genotypes are shown: S10 indicates the dominant allele associated with partial virulence while s10 corresponds to the recessive avirulent allele.

According to this population structure analysis, *Rpv12*-breaking strains are genetically similar to avirulent strains collected near them and do not form a distinct genetic group (Fig. 4). Thus, virulent strains do not belong to a unique lineage but rather emerged from local *P. viticola* populations. In addition, they displayed different lengths of deletion at the *AvrRpv12* locus (avr^Δ1-3^, Fig. 1D), which hints at the selection of independent mutation events in each location.

All strains with high aggressiveness on *Rpv10* were collected in the Upper Rhine Plain, a winegrowing region located around the French-German border. Intriguingly, they showed more genetic similarity to North American strains than the rest of the panel, as seen along the first PC (Fig. 4A). In particular, Pv1419 1, in which the *S-AvrRpv10* locus was identified (Fig. 2), was placed halfway between the North American group and avirulent strains from the Upper Rhine Plain. Genetic clustering revealed an admixture from the North American population in *Rpv10*-breaking strains, in variable proportions depending on the sample (17 to 57%) (Fig. 4B). This signal was not observed in avirulent strains from the same region. The balanced ancestry of Pv1419 1 suggests that it could be an early generation hybrid between populations, in accordance with its very high heterozygosity rate (1.41 heterozygous sites per kb versus 0.70 on average in non-admixed strains).

### The *S-AvrRpv10* locus coincides with an admixed genomic region

To understand if virulence towards *Rpv10* was linked to a potential North American admixture, we analyzed the levels of each ancestry along the chromosome 16 in admixed strains. Ancestral segments were defined using a subset of 3876 SNPs between the pure European and North American samples. Parent strain Pv1419 1 is heterozygous at the *S-AvrRpv10* locus (S10/s10) (Fig. 2D) and it showed a mix of the two ancestries along the entire chromosome (Fig.5). Descendants of Pv1419 1 had different ancestry profiles depending on their genotype. For example, cPv44 1 (S10/s10) presented a mixed ancestry while cPv43 1 (s10/s10) had a full European ancestry. In addition, an homozygous S10/S10 individual from the BC-1419 population presented a full American ancestry (bcPv44-04, Fig.5). The profile of wild strain Pv7219 1 was fully American along the locus except for the distal end where it was mixed. This pattern was consistent with the markers used to genotype the BC-1419 population (Data S5). Other *Rpv10*-breaking strains displayed either a mixed or a fully American ancestry.

Thus, the avirulent allele s10 is associated with a standard European genetic background, while the partially virulent allele S10 corresponds to an admixed region. This suggests that the adaptation to *Rpv10* occured through a genetic contribution from an extra-European population.

## Discussion

In this study, we successfully mapped two new loci in the *P. viticola* genome involved in the interaction with *Rpv10* and *Rpv12*, two key resistance factors of European grapevine breeding programs. Furthermore, sequencing a panel of strains representing the pathogen’s diversity provided insights into the dynamics of virulence emergence, both at the genomic and population levels.

### Dynamic regions foster rapid adaptation to resistances

The three major QTLs involved in resistance breakdown share many common features. They contain clusters of genes encoding secreted proteins with typical oomycete effector motifs (RXLR for *AvrRpv12*, dEER for *AvrRpv3*.*1* and *S-AvrRpv10*). These loci also display canonical or modified LWY domains, which participate in a modular structure. Closely related sequences tend to be physically close and the limit is often blurred between paralogous genes and additional copies. Analysis of different haplotypes confirmed that these loci are evolutionarily dynamic, with their effector repertoire varying between alleles. *AvrRpv3*.*1* and *AvrRpv12* are affected by deletions of different lengths that can comprise many genes in virulent alleles. The gene content fluctuates widely in the *S-AvrRpv10* locus, and large structural rearrangements are observed between parental haplotypes. In the *AvrRpv12* locus, CNV of RXLR genes occurs in both virulent and avirulent alleles, suggesting an evolution driven by repeated tandem duplications, possibly mediated by transposable elements and/or unequal crossovers (Dong et al, 2015; Fouché et al, 2018). Notably, this region exhibits a relatively high recombination rate (27.5 vs 18.0 cM/Mb on average in the genome). By contrast, large structural variations, such as those observed at the *S-AvrRpv10* locus, may hinder meiotic recombination when the divergence reaches a critical threshold. This could explain in part why we previously found that putative effector genes were enriched in poorly recombining regions of the *P. viticola* genome (Dvorak et al, 2025).

Overall, these observations are reminiscent of the genomic organization of other oomycete pathogens, in which effector genes tend to be located in highly dynamic regions (Haas et al, 2009; Qutob et al, 2009; Skiadas et al, 2024). This fast gene turnover in virulence-associated loci fosters a strong adaptative potential, in line with the two-speed genome model (Dong et al, 2015). Thus, effectors recognized by the plant can be lost, while new duplicated sequences can diverge to escape recognition or acquire new functions. The genetic diversity in terms of presence-absence variation of effectors is probably one of the key of the rapid adaptation of *P. viticola* populations to grapevine immune responses (Sánchez-Vallet et al, 2018).

### Parallel adaptation to *Rpv12* occurred in different regions by loss of avirulence

The loss of an Avr factor is one of the most common ways through which omycete plant pathogens gain virulence (Wang et al, 2019). In *P. viticola*, the absence of effector genes is necessary for the breakdown of *Rpv3*.*1* (Paineau et al, 2024). Our findings suggest that that the mode of adaptation to *Rpv12* is similar and probably linked to the loss of recognized effector(s). Both resistance breakdowns thus appear to be standard cases of ETI evasion. This is consistent with the dominance of avirulence, as a single Avr copy is sufficient to trigger HR and limit pathogen development.

Interestingly, we observed two types of virulence alleles at the *AvrRpv12* locus. Deletions (avr^Δ1-3^ alleles in Fig. 1D) are associated with the complete absence of HR. By contrast, the avr^ns^ allele produces a high level of sporulation but also visible necrosis in the area of pathogen growth. This “trailing necrosis” phenotype is commonly observed in partial resistance to downy mildew in *Arabidopsis thaliana* and it is generally interpreted as the result of a late or weaker immune response (Holub, 2001; Krasileva et al, 2011). Incomplete resistance to *Bremia lactucae* was also associated with a delayed HR in lettuce (Zhang et al, 2009). The analysis of the 412×2543 cross showed that avr^ns^ is recessive over the avirulent allele, but dominant over the fully virulent one (Avr > avr^ns^ > avr^Δ1^). Although we cannot completely rule out that minor loci may be involved, the aneuploid individual cPv462 1 confirmed that a strain carrying only avr^Δ1^ was fully virulent.

Three paralogous RXLR genes are consistently absent in deletion alleles, raising the question of whether the loss of several effectors is needed to escape Rpv12 recognition. The avr^ns^ allele contains a non-sense mutation in a single RXLR gene, P17g40, whose function is probably severely impaired by the premature termination. However, an incomplete immune response may still be triggered upon recognition of P17g38 and/or P17g39. Given that the introgressed *Rpv12* locus contains several NLR genes (Venuti et al, 2013; Frommer et al, 2023), each candidate effector might be recognized by a different R protein. Alternatively, a truncated P17g40 protein may still be sufficient to initiate the Rpv12-mediated HR, albeit with a delayed response due to the poor interaction. In the case of the flax rust effector AvrM, which directly interacts with its cognate resistance protein, recognition is based on the C-terminal domain, but proteins with large truncations outside of this domain induce weaker cell death (Catanzariti et al, 2010).

The occurrence of distinct virulence alleles in different winegrowing regions suggests that independent mutational events affecting *AvrRpv12* were selected. Moreover, wild *Rpv12*-breaking strains exhibited long ROH around the QTL, a notable observation given the high heterozygosity rate in *P. viticola* (Gobbin et al, 2006; Dussert et al, 2019). This pattern is consistent with a selective sweep at the locus, in line with the recent introduction of *Rpv12*-carrying varieties. At the regional level, the adaptive potential of *P. viticola* is thus sufficient to enable the rapid selection of Avr alleles escaping recognition, as was observed upon the deployment of *Rpv3*.*1* (Delmotte et al, 2014).

### A suppressor activity could play a role in virulence towards *Rpv10*

Unlike *Rpv3*.*1* and *Rpv12* breakdowns, virulence towards *Rpv10*-carrying plants is determined by a dominant locus. In diploid or dikaryotic pathogens, virulence is typically a recessive trait because a single avirulent allele is sufficient to trigger an immune response. However, seminal works on the flax rust pathosystem have described an Inhibitor (I) locus capable of preventing recognition of several Avr factors (Jones, 1988; Ellis et al, 2007). In several plant pathogens, certain effectors can indeed suppress the avirulence conferred by other genes. For example, the *Leptosphaeria maculans* effector AvrLm4-7 masks the presence of AvrLm3 which is normally recognized by Rlm3 (Plissonneau et al, 2016). A similar relationship exists between SvrPm3 and AvrPm3 in *Blumeria graminis* (Bourras et al, 2015), the first suppressing the Pm3-mediated recognition of the second. In the downy mildew pathogen *Hyaloperonospora arabidopsidis*, the secreted protein S-HAC1 suppresses the avirulence conferred by HAC1 (Woods-Tör et al, 2018). Thus, one of the putative effectors exclusive to the active *S-AvrRpv10* allele may effectively mask a hypothetical AvrRpv10 factor. The extensive repertoire of secreted proteins in *P. viticola* may have evolved, at least in part, to allow certain effectors to “protect” others by disrupting NLR-mediated immune responses. (Wu and Derevnina, 2023).

The interaction appears to be specific to the Rpv10-mediated response, as we found no effect of the QTL on plants carrying *Rpv3*.*1, Rpv12* or no resistance genes. Moreover, the donor strain Pv1419 1 is not aggressive towards *Rpv1* either (Paineau et al, 2022). However, the active *S-AvrRpv10* allele was not sufficient to totally prevent the HR in Pv1419 1 descendants, whereas little to no necrosis was observed with their parent. The wild strain Pv7219 1 also clearly induced necrosis despite being homozygous for this allele. Similarly, Heyman et al (2021) reported a German isolate highly aggressive towards *Rpv10* which triggered abundant necrosis. Several hypotheses may explain this incomplete suppression of the immune response. Undetected QTLs could contribute to the gain of virulence. As the *Rpv10* locus contains several NLR genes (Schwander et al, 2012), S-AvrRpv10 may mask only one protein among several recognized effectors. Alternatively, the suppression activity may vary between strains due to different expression levels, as observed for the *B. graminis* effector SvrPm3 (McNally et al, 2018).

### The breakdown of *Rpv10* was facilitated by admixture

In this study, we showed that adaptive admixture enabled the emergence of virulence towards *Rpv10* in Europe. This finding aligns with a broader pattern observed in several plant pathogens where introgression between divergent populations, and sometimes different (sub)species, has facilitated adaptation to new hosts and the evolution of distinct pathotypes (Menardo et al, 2016; Guo et al, 2022; Rahnama et al, 2023). A notable example is the breakdown of apple scab resistance gene *Rvi6* that occured after crabapple-associated *Venturia inaequalis* strains invaded orchards and later hybridized with the agricultural population, leading to the introgression of the virulent trait (Leroy et al, 2016; Michalecka et al, 2018). Here, the population structure analysis points to a recent North American origin of virulence towards *Rpv10* in *P. viticola*. This observation is intriguing as *Rpv10* was introgressed from the East-Asian species *V. amurensis*, and one could expect that adapted downy mildew strains would rather originate from the same region. However, as the center of origin of *P. viticola*, North America harbors an allelic diversity that is not fully represented in the rest of the world (Fontaine et al, 2021). Invasive species are generally characterized by a reduced genetic diversity in their new habitat, due to the genetic drift resulting from the small initial population size. This founder effect is clearly observed in *P. viticola* populations outside of North America (Fontaine et al, 2021). It is therefore plausible that recently introduced American genotypes included beneficial variants that facilitated the adaptation to *Rpv10*.

The detection of a new entry of the pathogen into Europe associated with resistance breakdown is particularly alarming. This reinforces the view that secondary introductions of already established invasive pathogens should be avoided as this can expand their allele reservoir and thus their adaptive potential (Dlugosch and Parker, 2008; Ahmed et al, 2012). In the case of grapevine downy mildew, pathogen movement from North America is especially concerning, because several *Plasmopara* species native to the region can infect wild and cultivated grapes (Rouxel et al, 2014; Mouafo-Tchinda et al, 2022).

## Conclusion

Overall, we demonstrated that virulence towards partial resistance genes was determined by major loci and we identified promising putative effectors involved in the interaction of *P. viticola* with Rpv10 and Rpv12. By integrating assessments of sporulation and necrosis, we were able to precisely characterize the extent of resistance breakdowns, whether partial or complete. We thus confirmed that gene-for-gene relationships are not restricted to complete resistances and that this model remains relevant in the context of quantitative interactions across various pathosystems (Jiquel et al, 2021; Langlands-Perry et al, 2023). We also identified a potential suppression activity targeting the Rpv10-mediated response, which could add another layer of complexity to the molecular interplay. Future functional assays will help decipher the role of candidate effectors, a challenging task for an obligate biotrophic pathogen like *P. viticola*. Co-expression of resistance and effector gene pairs would also require the cloning of Rpv10 and Rpv12, which has not yet been achieved to our knowledge.

Preserving the durability of resistances is critical for perennial plants such as grapevines, which are planted for decades with no possibility of crop or varietal rotation. Here, we show that linkage mapping and population genomics can be combined to understand the diverse pathways by which specialized plant pathogens acquire virulence. In *P. viticola*, parallel adaptation can occur independently in different established populations, while punctual admixture events can also contribute significantly to the emergence of virulence. Ensuring the long-term effectiveness of grapevine resistance will require accounting for the multiple trajectories of pathogen adaptation.

## Acknowledgments

We thank Julie Bourg, Carole Couture and Anne-Sophie Miclot (INRAE Bordeaux, France) for technical assistance, Frédéric Fabre for his advice on statistical analyses and Sabine Wiedemann-Merdinoglu (INRAE Colmar, France) for her insights on necrosis notation. We are grateful to the people who provided us with *P. viticola* strains: Pál Kozma (University of Pécs, Hungary), Lance Cadle-Davidson and Anna Underhill (Cornell University, Geneva, NY, United States), Odile Carisse (Horticultural Research and Development Centre, Agriculture and Agri-Food Canada), René Fuchs and Stefan Schumacher (State Institute of Viticulture and Oenology, Freiburg im Breisgau, Germany), Jochen Bogs, Birgit Eisenmann and Chantal Wingerter (Winecampus Neustadt, Germany), Elisa De Luca (Vivai Cooperativi Rauscedo, Rauscedo, Italy). We acknowledge the Genotoul bioinformatics platform (Toulouse, France) for providing computing and storage resources.

## Author contributions

ED, IDM and AC performed the experiments. MP and DC carried out the de novo genome assembly. ED, TD, PM, MFO and FD analyzed the data. FD, MFO and ED designed the study.

## Data availability

DNA sequences of *P. viticola* strains were submitted to NCBI SRA (project numbers indicated in Data S1). The genome assembly of Pv1419 1 and the phenotyping data produced in this study are available at https://doi.org/10.57745/KVUHYA.

## References

Ahmed S, de Labrouhe DT, Delmotte F (2012) Emerging virulence arising from hybridisation facilitated by multiple introductions of the sunflower downy mildew pathogen Plasmopara halstedii. Fungal Genetics and Biology 49(10):847–855. 10.1016/j.fgb.2012.06.012

Alexander DH, Lange K (2011) Enhancements to the ADMIXTURE algorithm for individual ancestry estimation. BMC Bioinformatics 12(1):246. 10.1186/1471-2105-12-246

Almagro Armenteros JJ, Tsirigos KD, Sønderby CK, Petersen TN, Winther O, Brunak S, von Heijne G, Nielsen H (2019) SignalP 5.0 improves signal peptide predictions using deep neural networks. Nat Biotechnol 37(4):420–423. 10.1038/s41587-019-0036-z

Arends D, Prins P, Jansen RC, Broman KW (2010) R/qtl: high-throughput multiple QTL mapping. Bioinformatics 26(23):2990–2992. 10.1093/bioinformatics/btq565

Bellin D, Peressotti E, Merdinoglu D, Wiedemann-Merdinoglu S, Adam-Blondon AF, Cipriani G, Morgante M, Testolin R, Di Gaspero G (2009) Resistance to Plasmopara viticola in grapevine ‘Bianca’ is controlled by a major dominant gene causing localised necrosis at the infection site. Theor Appl Genet 120(1):163–176. 10.1007/s00122-009-1167-2

Blum M, Waldner M, Gisi U (2010) A single point mutation in the novel PvCesA3 gene confers resistance to the carboxylic acid amide fungicide mandipropamid in Plasmopara viticola. Fungal Genetics and Biology 47(6):499–510. 10.1016/j.fgb.2010.02.009

Bourras S, McNally KE, Ben-David R, Parlange F, Roffler S, Praz CR, Oberhaensli S, Menardo F, Stirnweis D, Frenkel Z, et al (2015) Multiple Avirulence Loci and Allele-Specific Effector Recognition Control the Pm3 Race-Specific Resistance of Wheat to Powdery Mildew. The Plant Cell 27(10):2991–3012. 10.1105/tpc.15.00171

Boutemy LS, King SRF, Win J, Hughes RK, Clarke TA, Blumenschein TMA, Kamoun S, Banfield MJ (2011) Structures of Phytophthora RXLR Effector Proteins. Journal of Biological Chemistry 286(41):35834–35842. 10.1074/jbc.M111.262303

Browning BL, Tian X, Zhou Y, Browning SR (2021) Fast two-stage phasing of large-scale sequence data. The American Journal of Human Genetics 108(10):1880–1890. 10.1016/j.ajhg.2021.08.005

Casagrande K, Falginella L, Castellarin SD, Testolin R, Di Gaspero G (2011) Defence responses in Rpv3-dependent resistance to grapevine downy mildew. Planta 234(6):1097–1109. 10.1007/s00425-011-1461-5

Catanzariti AM, Dodds PN, Ve T, Kobe B, Ellis JG, Staskawicz BJ (2010) The AvrM Effector from Flax Rust Has a Structured C-Terminal Domain and Interacts Directly with the M Resistance Protein. MPMI 23(1):49–57. 10.1094/MPMI-23-1-0049

Chang CC, Chow CC, Tellier LC, Vattikuti S, Purcell SM, Lee JJ (2015) Second-generation PLINK: rising to the challenge of larger and richer datasets. GigaScience 4(1):s13742.–015–0047–8. 10.1186/s13742-015-0047-8

Combier M, Evangelisti E, Piron MC, Rengel D, Legrand L, Shenhav L, Bouchez O, Schornack S, Mestre P (2019) A secreted WY-domain-containing protein present in European isolates of the oomycete Plasmopara viticola induces cell death in grapevine and tobacco species. PLOS ONE 14(7):e0220184. 10.1371/journal.pone.0220184

Delmas CEL, Dussert Y, Delière L, Couture C, Mazet ID, Richart Cervera S, Delmotte F (2017) Soft selective sweeps in fungicide resistance evolution: recurrent mutations without fitness costs in grapevine downy mildew. Molecular Ecology 26(7):1936–1951. 10.1111/mec.14006

Delmotte F, Mestre P, Schneider C, Kassemeyer HH, Kozma P, Richart-Cervera S, Rouxel M, Delière L (2014) Rapid and multiregional adaptation to host partial resistance in a plant pathogenic oomycete: Evidence from European populations of Plasmopara viticola, the causal agent of grapevine downy mildew. Infection, Genetics and Evolution 27:500–508. 10.1016/j.meegid.2013.10.017

Dlugosch KM, Parker IM (2008) Founding events in species invasions: genetic variation, adaptive evolution, and the role of multiple introductions. Molecular Ecology 17(1):431–449. 10.1111/j.1365-294X.2007.03538.x

Dong S, Raffaele S, Kamoun S (2015) The two-speed genomes of filamentous pathogens: waltz with plants. Current Opinion in Genetics & Development 35:57–65. 10.1016/j.gde.2015.09.001

Dussert Y, Mazet ID, Couture C, Gouzy J, Piron MC, Kuchly C, Bouchez O, Rispe C, Mestre P, Delmotte F (2019) A High-Quality Grapevine Downy Mildew Genome Assembly Reveals Rapidly Evolving and Lineage-Specific Putative Host Adaptation Genes. Genome Biology and Evolution 11(3):954–969. 10.1093/gbe/evz048

Dvorak E, Mazet ID, Couture C, Delmotte F, Foulongne-Oriol M (2025) Recombination landscape and karyotypic variations revealed by linkage mapping in the grapevine downy mildew pathogen Plasmopara viticola. G3 Genes Genomes Genetics 15(1):jkae259. 10.1093/g3journal/jkae259

Ellis JG, Dodds PN, Lawrence GJ (2007) Flax Rust Resistance Gene Specificity is Based on Direct Resistance-Avirulence Protein Interactions. Annu Rev Phytopathol 45(1):289–306. 10.1146/annurev.phyto.45.062806.094331

Feechan A, Anderson C, Torregrosa L, Jermakow A, Mestre P, Wiedemann-Merdinoglu S, Merdinoglu D, Walker AR, Cadle-Davidson L, Reisch B, et al (2013) Genetic dissection of a TIR-NB-LRR locus from the wild North American grapevine species uscadinia rotundifolia identifies paralogous genes conferring resistance to major fungal and oomycete pathogens in cultivated grapevine. The Plant Journal 76(4):661–674. 10.1111/tpj.12327

Fletcher K, Shin OH, Clark KJ, Feng C, Putman AI, Correll JC, Klosterman SJ, Van Deynze A, Michelmore RW (2022) Ancestral Chromosomes for Family Peronosporaceae Inferred from a Telomere-to-Telomere Genome Assembly of Peronospora effusa. MPMI 35(6):450–463. 10.1094/MPMI-09-21-0227-R

Fontaine MC, Labbé F, Dussert Y, Delière L, Richart-Cervera S, Giraud T, Delmotte F (2021) Europe as a bridgehead in the worldwide invasion history of grapevine downy mildew, Plasmopara viticola. Current Biology 31(10):2155–2166.e4. 10.1016/j.cub.2021.03.009

Foria S, Copetti D, Eisenmann B, Magris G, Vidotto M, Scalabrin S, Testolin R, Cipriani G, Wiedemann-Merdinoglu S, Bogs J, et al (2020) Gene duplication and transposition of mobile elements drive evolution of the Rpv3 resistance locus in grapevine. The Plant Journal 101(3):529–542. 10.1111/tpj.14551

Fouché S, Plissonneau C, Croll D (2018) The birth and death of effectors in rapidly evolving filamentous pathogen genomes. Current Opinion in Microbiology 46:34–42. 10.1016/j.mib.2018.01.020

Frommer B, Müllner S, Holtgräwe D, Viehöver P, Huettel B, Töpfer R, Weisshaar B, Zyprian E (2023) Phased grapevine genome sequence of an Rpv12 carrier for biotechnological exploration of resistance to Plasmopara viticola. Front Plant Sci 14. 10.3389/fpls.2023.1180982

Fu Q, Chen T, Wang Y, Zhou H, Zhang K, Zheng R, Zhang Y, Liu R, Yin X, Liu G, et al (2024) Plasmopara viticola effector PvCRN20 represses the import of VvDEG5 into chloroplasts to suppress immunity in grapevine. New Phytologist 243(6):2311–2331. 10.1111/nph.20002

Gassmann W, Bhattacharjee S (2012) Effector-Triggered Immunity Signaling: From Gene-for-Gene Pathways to Protein-Protein Interaction Networks. MPMI 25(7):862–868. 10.1094/MPMI-01-12-0024-IA

Gobbin D, Rumbou A, Linde CC, Gessler C (2006) Population genetic structure of Plasmopara viticola after 125 years of colonization in European vineyards. Molecular Plant Pathology 7(6):519–531. 10.1111/j.1364-3703.2006.00357.x

Goddard TD, Huang CC, Meng EC, Pettersen EF, Couch GS, Morris JH, Ferrin TE (2018) UCSF ChimeraX: Meeting modern challenges in visualization and analysis. Protein Science 27(1):14–25. 10.1002/pro.3235

Guo Y, Betzen B, Salcedo A, He F, Bowden RL, Fellers JP, Jordan KW, Akhunova A, Rouse MN, Szabo LJ, et al (2022) Population genomics of Puccinia graminis f.sp. tritici highlights the role of admixture in the origin of virulent wheat rust races. Nat Commun 13(1):6287. 10.1038/s41467-022-34050-w

Haas BJ, Kamoun S, Zody MC, Jiang RHY, Handsaker RE, Cano LM, Grabherr M, Kodira CD, Raffaele S, Torto-Alalibo T, et al (2009) Genome sequence and analysis of the Irish potato famine pathogen Phytophthora infestans. Nature 461(7262):393–398. 10.1038/nature08358

He J, Ye W, Choi DS, Wu B, Zhai Y, Guo B, Duan S, Wang Y, Gan J, Ma W, et al (2019) Structural analysis of Phytophthora suppressor of RNA silencing 2 (PSR2) reveals a conserved modular fold contributing to virulence. Proceedings of the National Academy of Sciences 116(16):8054–8059. 10.1073/pnas.1819481116

Heyman L, Höfle R, Kicherer A, Trapp O, Ait Barka E, Töpfer R, Höfte M (2021) The Durability of Quantitative Host Resistance and Variability in Pathogen Virulence in the Interaction Between European Grapevine Cultivars and Plasmopara viticola. Front Agron 3. 10.3389/fagro.2021.684023

Holub EB (2001) The arms race is ancient history in Arabidopsis, the wildflower. Nat Rev Genet 2(7):516–527. 10.1038/35080508

Jiquel A, Gervais J, Geistodt-Kiener A, Delourme R, Gay EJ, Ollivier B, Fudal I, Faure S, Balesdent MH, Rouxel T (2021) A gene-for-gene interaction involving a ‘late’ effector contributes to quantitative resistance to the stem canker disease in Brassica napus. New Phytologist 231(4):1510–1524. 10.1111/nph.17292

Jombart T, Collins C (2015) Analysing genome-wide SNP data using adegenet 2.0.0. URL https://adegenet.r-forge.r-project.org/files/tutorial-genomics.pdf

Jones DA (1988) Genetic Properties of Inhibitor Genes in Flax Rust that Alter Avirulence to Virulence on Flax. Phytopathology 78(3):342. 10.1094/Phyto-78-342

Jumper J, Evans R, Pritzel A, Green T, Figurnov M, Ronneberger O, Tunyasuvunakool K, Bates R, Žídek A, Potapenko A, et al (2021) Highly accurate protein structure prediction with AlphaFold. Nature 596(7873):583–589. 10.1038/s41586-021-03819-2

Juraschek L, Matera C, Steiner U, Oerke EC (2022) Pathogenesis of Plasmopara viticola Depending on Resistance Mediated by Rpv3 1, and Rpv10 and Rpv3 3, and by the Vitality of Leaf Tissue. Phytopathology® 112(7):1486–1499. 10.1094/PHYTO-10-21-0415-R

Kelley LA, Mezulis S, Yates CM, Wass MN, Sternberg MJE (2015) The Phyre2 web portal for protein modeling, prediction and analysis. Nat Protoc 10(6):845–858. 10.1038/nprot.2015.053

Koeck M, Hardham AR, Dodds PN (2011) The role of effectors of biotrophic and hemibiotrophic fungi in infection. Cellular Microbiology 13(12):1849–1857. 10.1111/j.1462-5822.2011.01665.x

Krasileva KV, Zheng C, Leonelli L, Goritschnig S, Dahlbeck D, Staskawicz BJ (2011) Global Analysis of Arabidopsis/Downy Mildew Interactions Reveals Prevalence of Incomplete Resistance and Rapid Evolution of Pathogen Recognition. PLOS ONE 6(12):e28765. 10.1371/journal.pone.0028765

Langlands-Perry C, Pitarch A, Lapalu N, Cuenin M, Bergez C, Noly A, Amezrou R, Gélisse S, Barrachina C, Parrinello H, et al (2023) Quantitative and qualitative plant-pathogen interactions call upon similar pathogenicity genes with a spectrum of effects. Front Plant Sci 14. 10.3389/fpls.2023.1128546

Lenth RV (2024) emmeans: Estimated Marginal Means, aka Least-Squares Means. URL https://CRAN.R-project.org/package=emmeans

Leroy T, Caffier V, Celton JM, Anger N, Durel CE, Lemaire C, Le Cam B (2016) When virulence originates from nonagricultural hosts: evolutionary and epidemiological consequences of introgressions following secondary contacts in Venturia inaequalis. New Phytologist 210(4):1443–1452. 10.1111/nph.13873

Lozano-Isla F (2024) inti: Tools and Statistical Procedures in Plant Science. URL https://CRAN.R-project.org/package=inti

Maddalena G, Delmotte F, Bianco PA, De Lorenzis G, Toffolatti SL (2020) Genetic structure of Italian population of the grapevine downy mildew agent, Plasmopara viticola. Annals of Applied Biology 176(3):257–267. 10.1111/aab.12567

Matson MEH, Liang Q, Lonardi S, Judelson HS (2022) Karyotype variation, spontaneous genome rearrangements affecting chemical insensitivity, and expression level polymorphisms in the plant pathogen Phytophthora infestans revealed using its first chromosome-scale assembly. PLOS Pathogens 18(10):e1010869. 10.1371/journal.ppat.1010869

McDonald BA, Linde C (2002) Pathogen population genetics, evolutionary potential, and durable resistance. Annual Review of Phytopathology 40(Volume 40, 2002):349–379. 10.1146/annurev.phyto.40.120501.101443

McNally KE, Menardo F, Lüthi L, Praz CR, Müller MC, Kunz L, Ben-David R, Chandrasekhar K, Dinoor A, Cowger C, et al (2018) Distinct domains of the AVRPM3A2/F2 avirulence protein from wheat powdery mildew are involved in immune receptor recognition and putative effector function. New Phytologist 218(2):681–695. 10.1111/nph.15026

Menardo F, Praz CR, Wyder S, Ben-David R, Bourras S, Matsumae H, McNally KE, Parlange F, Riba A, Roffler S, et al (2016) Hybridization of powdery mildew strains gives rise to pathogens on novel agricultural crop species. Nat Genet 48(2):201–205. 10.1038/ng.3485

Michalecka M, Masny S, Leroy T, Pu-lawska J (2018) Population structure of Venturia inaequalis, a causal agent of apple scab, in response to heterogeneous apple tree cultivation. BMC Evol Biol 18(1):5. 10.1186/s12862-018-1122-4

Minio A, Cochetel N, Vondras AM, Massonnet M, Cantu D (2022) Assembly of complete diploid-phased chromosomes from draft genome sequences. G3 Genes textbackslashtextbarGenomes textbackslashtextbarGenetics 12(8):jkac143. 10.1093/g3journal/jkac143

Mirdita M, Schütze K, Moriwaki Y, Heo L, Ovchinnikov S, Steinegger M (2022) ColabFold: making protein folding accessible to all. Nat Methods 19(6):679–682. 10.1038/s41592-022-01488-1

Mouafo-Tchinda RA, Fall ML, Beaulieu C, Carisse O (2022) Competition Between Plasmopara viticola Clade riparia and Clade aestivalis: A Race to Lead Grape Downy Mildew Epidemics. Plant Disease 106(11):2866–2875. 10.1094/PDIS-11-21-2465-RE

Möller E, Bahnweg G, Sandermann H, Geiger H (1992) A simple and efficient protocol for isolation of high molecular weight DNA from filamentous fungi, fruit bodies, and infected plant tissues. Nucleic Acids Research 20(22):6115–6116. 10.1093/nar/20.22.6115

Paineau M, Mazet ID, Wiedemann-Merdinoglu S, Fabre F, Delmotte F (2022) The Characterization of Pathotypes in Grapevine Downy Mildew Provides Insights into the Breakdown of Rpv3, Rpv10, and Rpv12 Factors in Grapevines. Phytopathology® 112(11):2329–2340. 10.1094/PHYTO-11-21-0458-R

Paineau M, Minio A, Mestre P, Fabre F, Mazet ID, Couture C, Legeai F, Dumartinet T, Cantu D, Delmotte F (2024) Multiple deletions of candidate effector genes lead to the breakdown of partial grapevine resistance to downy mildew. New Phytologist 243(4):1490–1505. 10.1111/nph.19861

Petit-Houdenot Y, Fudal I (2017) Complex Interactions between Fungal Avirulence Genes and Their Corresponding Plant Resistance Genes and Consequences for Disease Resistance Management. Front Plant Sci 8. 10.3389/fpls.2017.01072

Plissonneau C, Daverdin G, Ollivier B, Blaise F, Degrave A, Fudal I, Rouxel T, Balesdent MH (2016) A game of hide and seek between avirulence genes AvrLm4-7 and AvrLm3 in Leptosphaeria maculans. New Phytologist 209(4):1613–1624. 10.1111/nph.13736

Possamai T, Wiedemann-Merdinoglu S (2022) Phenotyping for QTL identification: A case study of resistance to Plasmopara viticola and Erysiphe necator in grapevine. Front Plant Sci 13. 10.3389/fpls.2022.930954

Qutob D, Tedman-Jones J, Dong S, Kuflu K, Pham H, Wang Y, Dou D, Kale SD, Arredondo FD, Tyler BM, et al (2009) Copy Number Variation and Transcriptional Polymorphisms of Phytophthora sojae RXLR Effector Genes Avr1a and Avr3a. PLOS ONE 4(4):e5066. 10.1371/journal.pone.0005066

Rahnama M, Condon B, Ascari JP, Dupuis JR, Del Ponte EM, Pedley KF, Martinez S, Valent B, Farman ML (2023) Recent co-evolution of two pandemic plant diseases in a multi-hybrid swarm. Nat Ecol Evol 7(12):2055–2066. 10.1038/s41559-023-02237-z

Rouxel M, Mestre P, Baudoin A, Carisse O, Delière L, Ellis MA, Gadoury D, Lu J, Nita M, Richard-Cervera S, et al (2014) Geographic Distribution of Cryptic Species of Plasmopara viticola Causing Downy Mildew on Wild and Cultivated Grape in Eastern North America. Phytopathology® 104(7):692–701. 10.1094/PHYTO-08-13-0225-R

Rouxel T, Balesdent MH (2010) Avirulence Genes. In: eLS. John Wiley & Sons, Ltd, 10.1002/9780470015902.a0021267

Salter-Townshend M, Myers S (2019) Fine-Scale Inference of Ancestry Segments Without Prior Knowledge of Admixing Groups. Genetics 212(3):869–889. 10.1534/genetics.119.302139

Schwander F, Eibach R, Fechter I, Hausmann L, Zyprian E, Töpfer R (2012) Rpv10: a new locus from the Asian Vitis gene pool for pyramiding downy mildew resistance loci in grapevine. Theor Appl Genet 124(1):163–176. 10.1007/s00122-011-1695-4

Skiadas P, Vidal SR, Dommisse J, Mendel MN, Elberse J, Ackerveken GVd, Jonge Rd, Seidl MF (2024) Pangenome graph analysis reveals extensive effector copy-number variation in spinach downy mildew. PLOS Genetics 20(10):e1011452. 10.1371/journal.pgen.1011452

Stassen JH, Van den Ackerveken G (2011) How do oomycete effectors interfere with plant life? Current Opinion in Plant Biology 14(4):407–414. 10.1016/j.pbi.2011.05.002

Śanchez-Vallet A, Fouché S, Fudal I, Hartmann FE, Soyer JL, Tellier A, Croll D (2018) The Genome Biology of Effector Gene Evolution in Filamentous Plant Pathogens. Annual Review of Phytopathology 56(Volume 56, 2018):21–40. 10.1146/annurev-phyto-080516-035303

Venuti S, Copetti D, Foria S, Falginella L, Hoffmann S, Bellin D, Cindrić P, Kozma P, Scalabrin S, Morgante M, et al (2013) Historical Introgression of the Downy Mildew Resistance Gene Rpv12 from the Asian Species Vitis amurensis into Grapevine Varieties. PLOS ONE 8(4):e61228. 10.1371/journal.pone.0061228

Wang Y, Tyler BM, Wang Y (2019) Defense and Counterdefense During Plant-Pathogenic Oomycete Infection. Annual Review of Microbiology 73(Volume 73, 2019):667–696. 10.1146/annurev-micro-020518-120022

Wingerter C, Eisenmann B, Weber P, Dry I, Bogs J (2021) Grapevine Rpv3-, Rpv10- and Rpv12-mediated defense responses against Plasmopara viticola and the impact of their deployment on fungicide use in viticulture. BMC Plant Biol 21(1):470. 10.1186/s12870-021-03228-7

Woods-Tör A, Studholme DJ, Cevik V, Telli O, Holub EB, Tör M (2018) A Suppressor/Avirulence Gene Combination in Hyaloperonospora arabidopsidis Determines Race Specificity in Arabidopsis thaliana. Front Plant Sci 9. 10.3389/fpls.2018.00265

Wu Ch, Derevnina L (2023) The battle within: How pathogen effectors suppress NLR-mediated immunity. Current Opinion in Plant Biology 74:102396. 10.1016/j.pbi.2023.102396

Xiang J, Li X, Wu J, Yin L, Zhang Y, Lu J (2016) Studying the Mechanism of Plasmopara viticola RxLR Effectors on Suppressing Plant Immunity. Front Microbiol 7:709. 10.3389/fmicb.2016.00709

Zhang NW, Pelgrom K, Niks RE, Visser RGF, Jeuken MJW (2009) Three Combined Quantitative Trait Loci from Nonhost Lactuca saligna Are Sufficient to Provide Complete Resistance of Lettuce Against Bremia lactucae. MPMI 22(9):1160– 1168. 10.1094/MPMI-22-9-1160

