## Supplemental File for "Parallel adaptation and admixture drive the evolution of virulence in the grapevine downy mildew pathogen"

This file includes:

**Methods S1** Chromosome-level assembly of strain Pv1419_1

**Methods S2** Genotyping of the backcross population by amplicon length polymorphism

**Methods S3** Variant calling and filtration

**Fig. S1** Notation of necrosis score on grapevine leaf discs infected by *P. viticola*

**Fig. S2** QTL mapping of the two *P. viticola* F1 progenies inoculated on susceptible or resistant grapevine cultivars.

**Fig. S3** Confirmation of the *AvrRpv3.1* locus in a *P. viticola* biparental population.

**Fig. S4** Maximum likelihood phylogenetic tree of RXLR protein sequences around the *AvrRpv12* locus

**Fig. S5** Coverage at the *AvrRpv12* locus in avirulent and virulent *P. viticola* strains

**Fig. S6** Runs of Homozygosity around the *AvrRpv12* locus in virulent *P. viticola* strains

**Fig. S7** Predicted tertiary structure of an AvrRpv12 candidate protein

**Table S1** Assembly statistics of the Pv1419_1 genome

**Table S2** PCR-based amplicon length polymorphism markers used to determine *S-AvrRpv10* alleles

**Methods S1** Chromosome-level assembly of strain Pv1419_1, quality assessment and annotation

HiFi reads of Pv1419_1 were assembled using a two-step procedure combining Hifiasm v0.16.1-r37412 (Cheng et al, 2021) and the HaploSync tool suite v1.0 (Minio et al, 2022). To generate the most contiguous and high-quality draft genome assembly, multiple parameter combinations were tested in Hifiasm, optimizing for haplotype size, contig length, and fragmentation. A total of 252 diploid assemblies were generated to identify the best configuration. The ten most promising assemblies were then screened for contamination. To remove contaminants, genetic markers developed in Dvorak et al (2025) were used to identify contigs belonging to the *P. viticola* genome. Contigs lacking these markers were aligned against the RefSeq genome (O'Leary et al, 2016) and classified using MEGAN6 (Huson et al, 2016) to identify contigs within the oomycete class. Verified oomycete contigs were incorporated into the *P. viticola* assembly, and contigs with no assignment were also retained. The assemblies with the highest quality were selected for the second step. Next, scaffolded, phased, chromosome-scale pseudomolecules were constructed for both haplotypes using the *P. viticola* genetic map (Dvorak et al, 2025) and HaploSync tool suite v1.0 (Table S1). Two iterations of HaploSplit and two iterations of HaploFill were performed to reduce fragmentation and finalize the assembly.

Genome completeness was evaluated using BUSCO v5 (Simão et al, 2015) with the alveolata_db10 (171 orthologs) and stramenopiles_odb10 (100 orthologs) datasets (Table S1). Analyses were performed separately for haplotypes 1 and 2 as well as for the unplaced contigs. Telomere repeat units were examined using TIDK v.0.2.63 (Brown et al, 2025). The analysis was conducted with tidk explore, targeting the terminal 1% of each chromosome and considering repeat units between 5 and 12 base pairs in length, with a minimum of two consecutive repetitions. The canonical telomeric repeat sequence TTTAGGG (Fulnečková et al, 2013) was subsequently identified using tidk search. Visualization revealed a nearly complete assembly from telomere to telomere. Telomeric repeats were absent at one end of chromosomes 5 and 17 in both haplotype 1 and 2, as well as chromosome 14 in haplotype 2.

The annotation from the most recent assembly of reference strain Pv221_1 (available at https://doi.org/10.57745/MXJWZS) was transferred to the two haplotype assemblies of Pv1419_1 using liftoff v1.6.2 (Shumate and Salzberg, 2021). Additionally, an open reading frame (ORF) search was performed along the QTL detected in this strain to reveal single-exon coding sequences that were absent from the reference genome. ORFs with a minimum length of 200 pb were detected using r/ORFik v1.24 (Tjeldnes et al, 2021). Amino acid sequences were extracted and submitted to InterProScan to identify protein domains (Jones et al, 2014). Sequences displaying transposon-associated domains were discarded. The remaining protein sequences were predicted to be secreted by detecting signal peptides with SignalP v5.0 (Almagro Armenteros et al, 2019).

**Methods S2** Genotyping of the backcross population by amplicon length polymorphism

Specific primers were designed to amplify DNA segments encompassing three indels, thus generating different amplicon lengths between alleles : 430 vs 766 pb for the first marker, 240 vs 199 pb for the second, and 153 versus 119 pb for the third. PCR amplifications were performed in a 20 µL reaction volume consisting of 8.2 µL of nuclease-free water, 10 µL of Platinum™ II Hot-Start PCR Master Mix , 0.4 µL of each primer at a concentration of 10 µM, and 1 µL of DNA template. The reactions were performed in a Applied Biosystems™ VeritiPro™ Thermal Cycler with the following program: 120 s at 94°C for initial denaturation, followed by 40 cycles of denaturation (30 s at 94°C), annealing (15 s at 55 or 60°C depending on the primers) and extension (30 s at 68°C). PCR products were visualized by electrophoresis on agarose gels. Primer sequences and annealing temperatures are indicated in Table S2.

**Methods S3** Mapping, variant calling and filtration

We used a pipeline implemented in RattleSNP (https://rattlesnp.readthedocs.io/) to parallelize read mapping and variant calling. Mapping on the Pv221_1 primary haplotype assembly was done with bwa-mem v0.7.18 (Li, 2013). Bam files of different libraries of the same sample were merged using samtools v1.18 (Li et al, 2009). Variant calling was performed using GATK v4.2.6 (McKenna et al, 2010). PCR duplicate reads were removed with the command Picard MarkDuplicates. Genotypes were called using GATK HaplotypeCaller with default parameters. Variants in repetitive regions were filtered out with bedtools v2.30 (Quinlan and Hall, 2010). High-confidence SNPs were retained using bcftools v1.17. Biallelic SNPs were the only sites considered. Sites with a depth twice as high as the average individual coverage were set to missing as they could be the result of false heterozygosity due to non-annotated transposable elements (TE) or copy number variation (CNV). SNPs with a depth inferior to 5 in a sample were set to missing. Finally, variants missing in more than 10% of the population were removed.

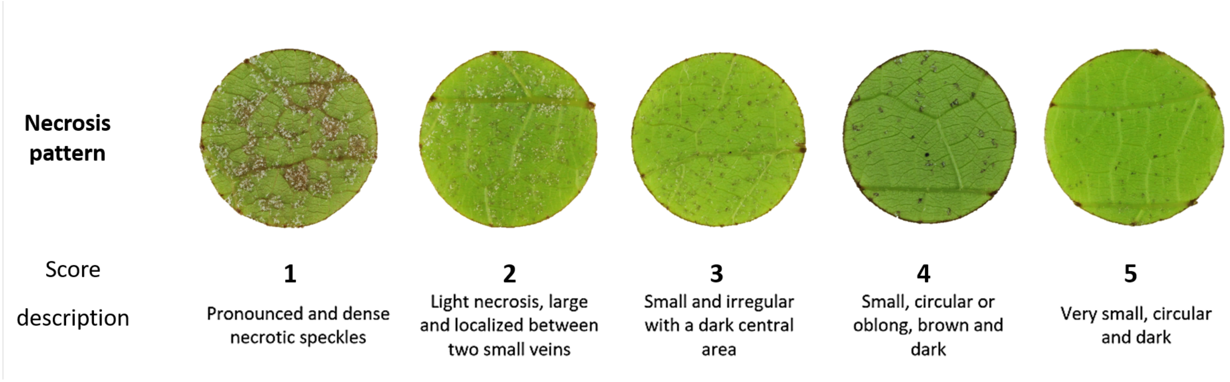


**Fig. S1**: **Notation of necrosis score on grapevine leaf discs infected by *P. viticola*.** Necrosis patterns on leaf discs were evaluated at 6 dpi by assessing their size, shape and color. Higher necrosis scores correspond to more efficient immune responses. Adapted from the scale presented in Paineau et al (2022).

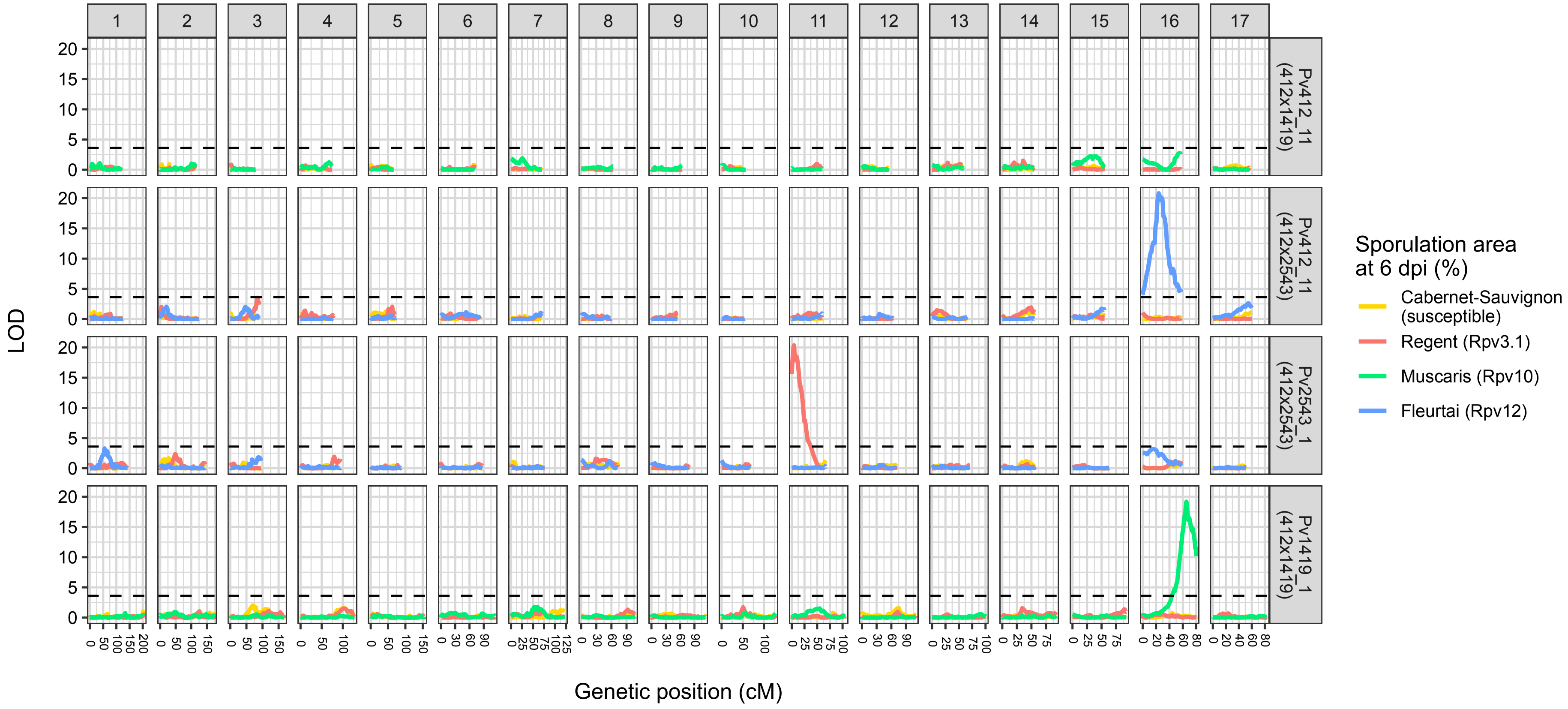


**Fig. S2**: **Simple interval QTL mapping of the two *P. viticola* F1 progenies inoculated on susceptible or resistant grapevine cultivars.** One map was obtained for each parent of each cross, as presented in Dvorak et al (2025). The detection of a QTL in one of the parental maps indicate a phenotypic difference in the progeny depending on which marker alleles were transmitted by the parent. The black dashed line indicate a LOD significance level of 3.2 which was the lowest threshold value determined across the different QTL mappings ($\alpha = 0.05$). LOD values were computed based on the percentage of sporulation area at 6 dpi.

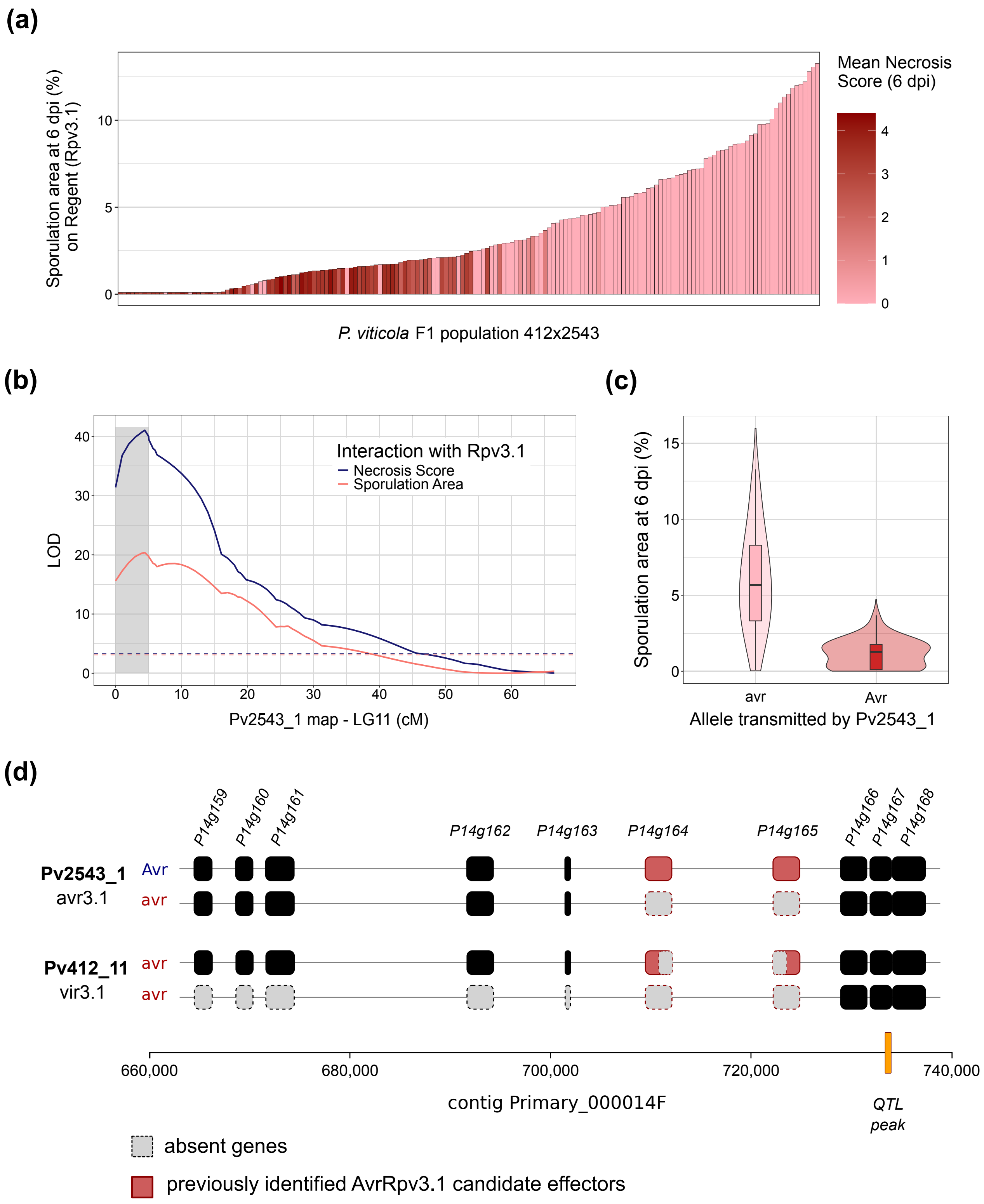


**Fig. S3**: **Confirmation of the AvrRpv3.1 locus in a *P. viticola* biparental population. (a)** Phenotypes distribution in the 412x2543 F1 progeny (N=162) on cv. 'Regent' (Rpv3.1). **(b)** QTL mapping of Rpv3.1-breakdown in the linkage map of the avirulent parent Pv2543_1. The gray area indicates the credibility interval of the QTL. Dashed lines indicate the LOD significance thresholds determined using 1000 permutations. Results on other linkage groups and other cultivars are available in Fig. S2. **(c)** Distribution of the sporulation area on Rpv3.1 depending on the inherited allele at the QTL. Horizontal lines in the boxplots signal the 25$^{th}$, 50$^{th}$ and 75$^{th}$ percentiles. **(d)** Allelic configurations of the parent strains in the region previously identified by GWAS (Paineau et al, 2024). The marker corresponding to the peak of the QTL in the present study is indicated by an orange bar on the scale. The allele associated with avirulence corresponds to the non-deleted Pv2543_1 haplotype (named Avr on the left). The secreted proteins P14g164 and P14g165 (colored in red) are totally or partially deleted in the virulent haplotypes.

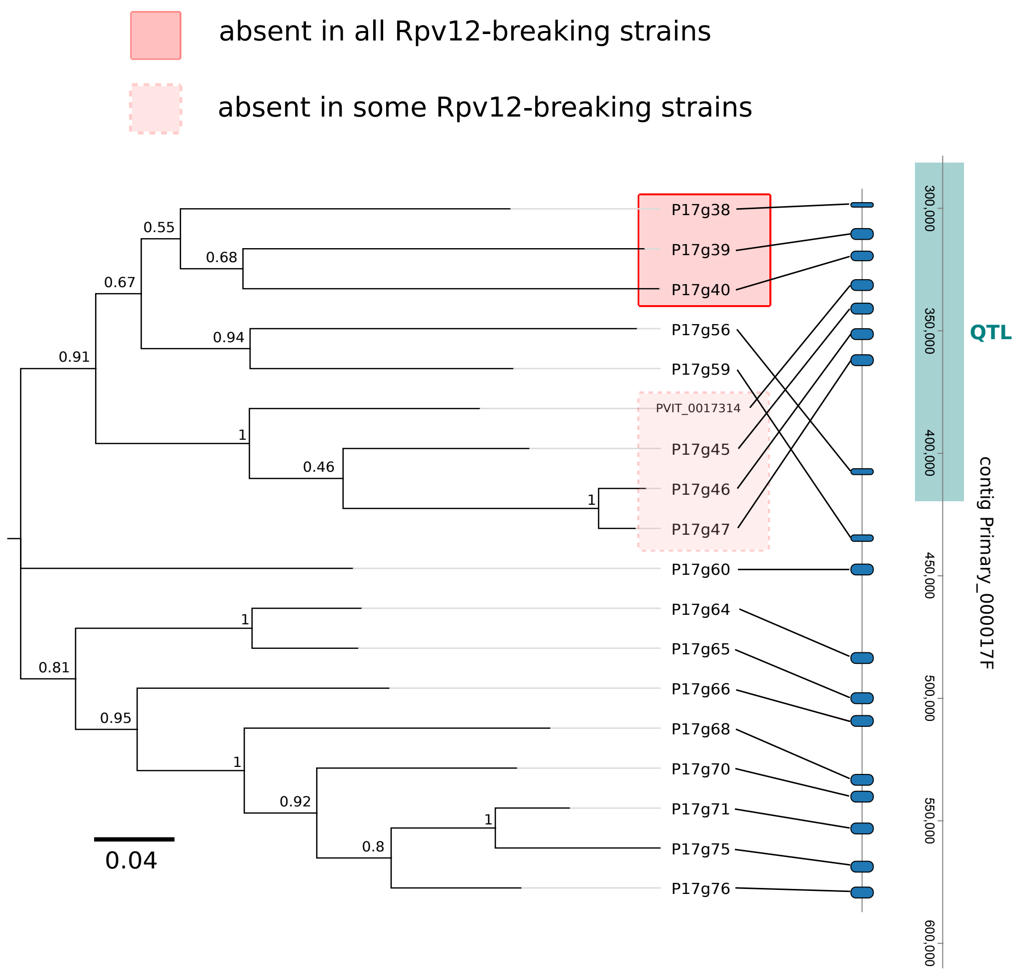


**Fig. S4**: **Maximum likelihood phylogenetic tree of RXLR protein sequences around the AvrRpv12 locus.** Sequence clustering using a 50% identity threshold revealed that RXLR genes in the QTL belonged to an extended family exclusively located in this genomic region. Bootstrap support values obtained from 1000 replicates are indicated for each node. RXLR genes absent in all or some Rpv12-breaking strains are highlighted in red boxes. The credibility interval of the AvrRpv12 QTL on contig Primary_000017F is indicated in turquoise.

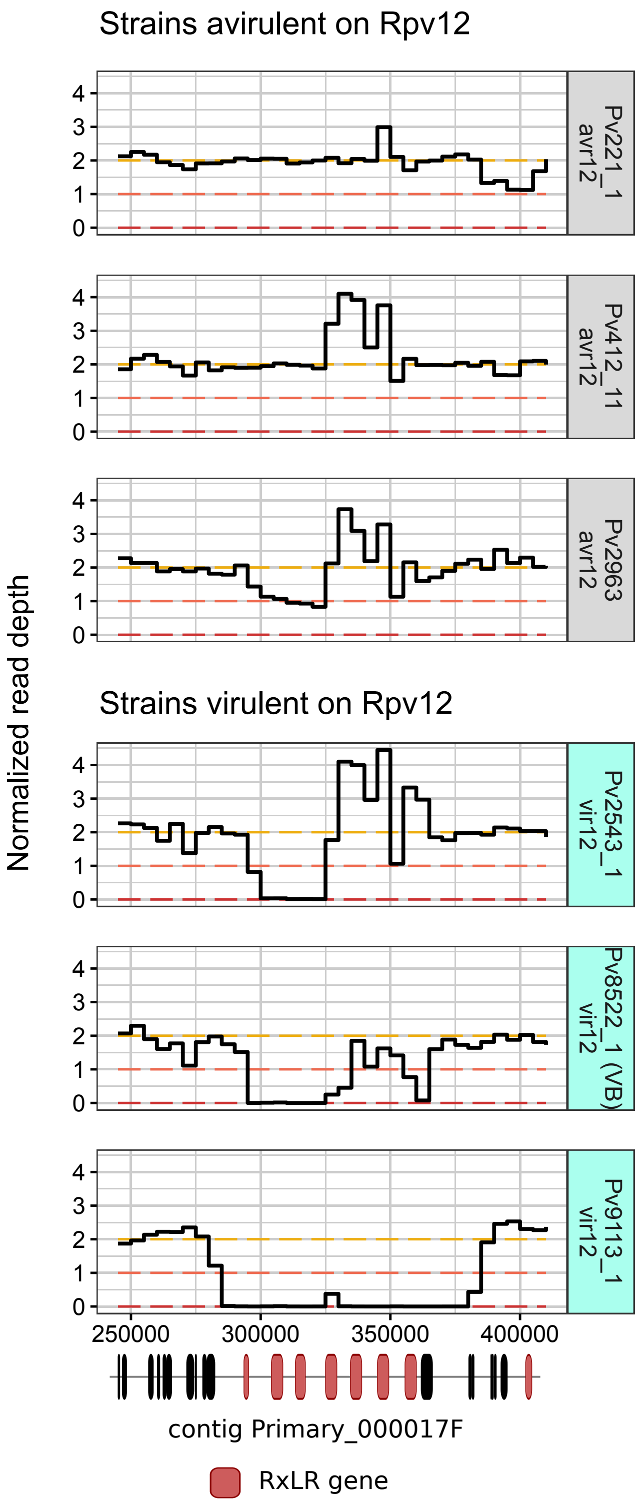


**Fig. S5**: **Depth of sequencing at the AvrRpv12 locus in avirulent and virulent *P. viticola* strains.** Normalized read depth along the QTL. Copy number is indicated on the y-axis and calculated by 5-kb windows. At the bottom, coding sequences are indicated in black, or in red for RXLR genes. Reference avirulent strain Pv221_1 is at the top. Blue boxes signal strains virulent on Rpv12. Some strains present higher coverage for the fourth and fifth RXLR genes, suggesting they possess additional copies. Note the hemizygous profile of strain Pv2963, which was collected from the same plot as Pv2543_1 but is avirulent.

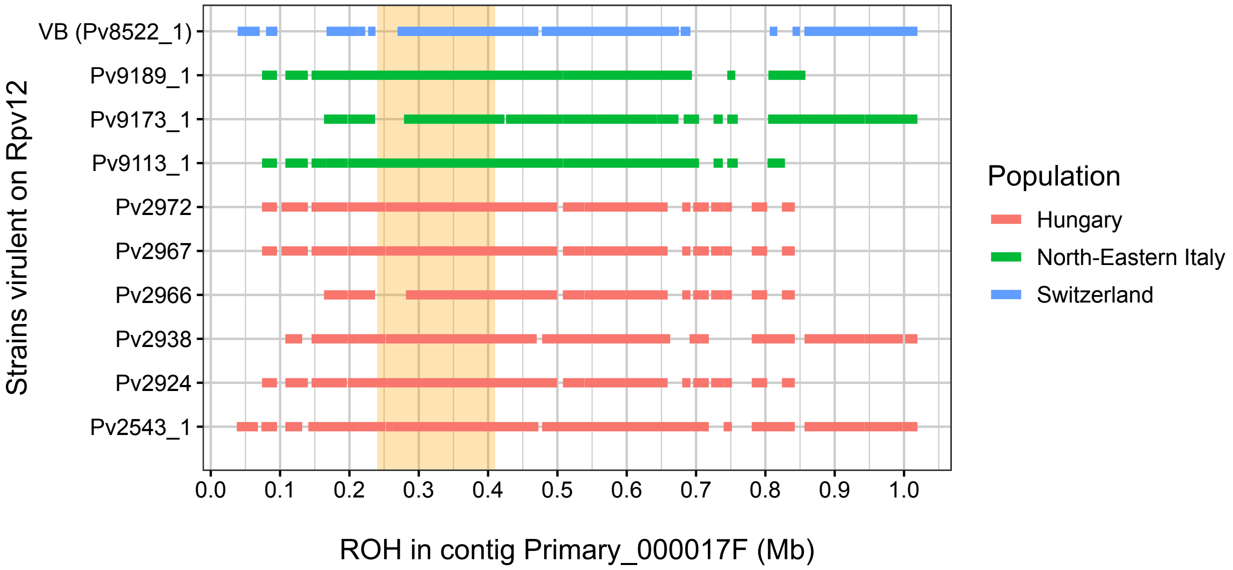


**Fig. S6**: **Runs of Homozygosity (ROH) around the AvrRpv12 locus in virulent *P. viticola* strains.** Lines indicate uninterrupted homozygous segments. They are colored according to the geographical origin of corresponding strains. The VB strain was presented in Wingerter et al (2021). Plot made using r/detectRUNS v0.9.6.

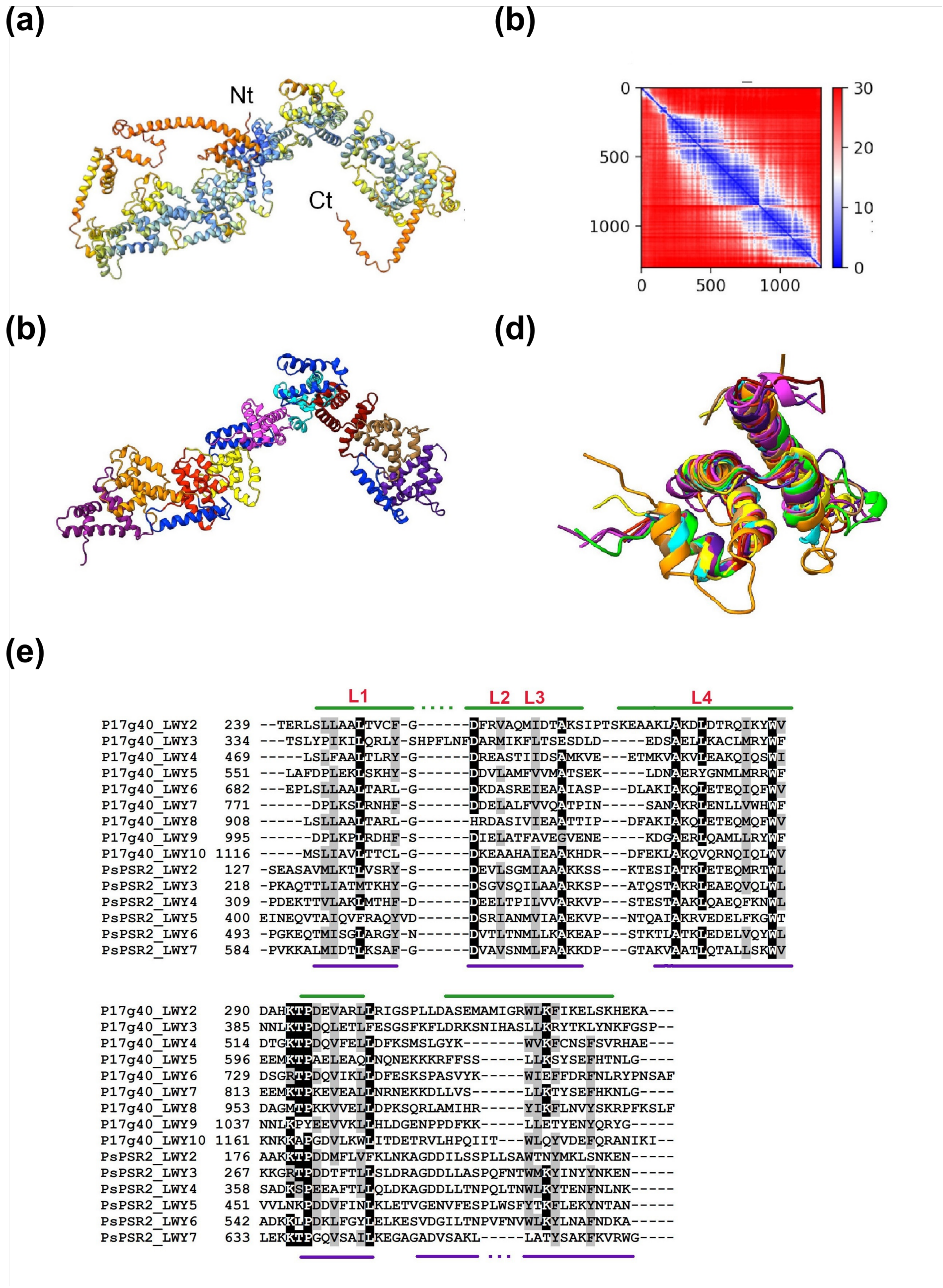


**Fig. S7**: **Predicted tertiary structure of an AvrRpv12 candidate protein.** P17g40 code for an RXLR protein composed of several modules of the LWY domain. **(a)** Complete AlphaFold-predicted structure colored by predicted local distance difference test (pLDDT) (red: low confidence, blue: high confidence). The N terminus and C terminus of the molecule are indicated by 'Nter' and 'Cter'. **(b)** Predicted aligned error (pAE) of the relative position of residues along the protein sequence. **(c)** Predicted structures with the 9 complete LWY modules highlighted in different colors. Sequences linking the different modules are shown in blue. Poorly predicted N- and C-terminal parts were trimmed for visual clarity. **(d)** Superimposition of all LWY domains of P17g40, using the third domain from the oomycete effector PsPSR2 as a reference. **(e)** Alignment of the P17g40 and the PsPSR2 sequences, skipping the first module that is shorter than the others in both proteins (He et al, 2019). Green lines at the top of the alignment indicate alpha-helices sequences for P17g40, and purple lines at the bottom indicate those of PsPSR2. Conserved leucine residues contributing to the fold are highlighted in red. Black background indicate identity and gray background similarity (70% cutoff).

**Tab. S1**: Assembly statistics of the Pv1419_1 genome

| HiFiasm assembly | Haplotype 1 | Haplotype 2 |
|---|---|---|
| Total contig length (bp) | 112,759,349 | 115,248,436 |
| N. of contigs | 88 | 69 |
| Contig N50 Length (bp) | 5,075,331 | 5,040,760 |
| Contig N50 index | 8 | 9 |
| Maximum contig length (bp) | 14,978,608 | 12,725,610 |

| HaploSync pseudomolecule reconstruction | Haplotype 1 | Haplotype 2 | Unplaced contigs |
|---|---|---|---|
| Total pseudomolecule length (bp) | 109,235,593 | 106,290,128 | 12,500,064 |
| N. of pseudomolecules | 17 | 17 | 105 |
| N. of pseudomolecules with 2 terminal telomeric repeats | 15 | 14 | - |
| N. of pseudomolecules with only 1 terminal telomeric repeat | 2 | 3 | - |
| GC content (%) | 45 | 45 | 44 |
| Complete BUSCO (%) (alveolata_odb10) | 95.9 | 95.9 | 0.6 |
| Complete BUSCO (%)(stramenopiles_odb10) | 100.0 | 100.0 | 0 |

**Tab. S2**: PCR-based amplicon length polymorphism markers used to determine S-AvrRpv10 alleles.

| Marker | Primers | Annealing tempera-ture (°C) | Amplicon length HAP1 | Amplicon length HAP2 | Position Pv221 Primary assembly | Position Pv1419 HAP1 (Chr16) | Position Pv1419 HAP2 (Chr16) |
|---|---|---|---|---|---|---|---|
| QTL-S10-MK1 | F: GAAGGAGAAGGTGGCAGCTG<br>R: GGTACATCGTTTTCTGCGCC | 60 | 430 | 766 | P62 222970-223400 | 3946627-3947057 | 4137410-4138176 |
| QTL-S10-MK2 | F: GTCGGCTGGGGGAATAAGTA<br>R: TTTCGTGGGATGCTCCAACTT | 60 | 240 | 199 | P62 334960-335200 | 4059313-4059553 | 4258061-4258260 |
| QTL-S10-MK3 | F: GTCGAAAGGAAACGTGACCC<br>R: GAGTGGCTCTCGCAAAACTT | 55 | 153 | 119 | P44 344935-345088 | 4405043-4405196 | 4708160-4708279 |

### Supplemental references


Almagro Armenteros JJ, Tsirigos KD, Sønderby CK, Petersen TN, Winther O, Brunak S, von Heijne G, Nielsen H (2019) SignalP 5.0 improves signal peptide predictions using deep neural networks. Nat Biotechnol 37(4):420–423. https://doi.org/10.1038/s41587-019-0036-z

Brown MR, Manuel Gonzalez de La Rosa P, Blaxter M (2025) tidk: a toolkit to rapidly identify telomeric repeats from genomic datasets. Bioinformatics 41(2):btaf049. https://doi.org/10.1093/bioinformatics/btaf049

Cheng H, Concepcion GT, Feng X, Zhang H, Li H (2021) Haplotype-resolved de novo assembly using phased assembly graphs with hifiasm. Nat Methods 18(2):170–175. https://doi.org/10.1038/s41592-020-01056-5

Dvorak E, Mazet ID, Couture C, Delmotte F, Foulongne-Oriol M (2025) Recombination landscape and karyotypic variations revealed by linkage mapping in the grapevine downy mildew pathogen Plasmopara viticola. G3 Genes|Genomes|Genetics 15(1):jkae259. https://doi.org/10.1093/g3journal/jkae259

Fulnečková J, Ševčíková T, Fajkus J, Lukešová A, Lukeš M, Vlček Lang BF, Kim E, Eliáš M, Sýkorová E (2013) A Broad Phylogenetic Survey Unveils the Diversity and Evolution of Telomeres in Eukaryotes. Genome Biology and Evolution 5(3):468–483. https://doi.org/10.1093/gbe/evt019

He J, Ye W, Choi DS, Wu B, Zhai Y, Guo B, Duan S, Wang Y, Gan J, Ma W, et al (2019) Structural analysis of Phytophthora suppressor of RNA silencing 2 (PSR2) reveals a conserved modular fold contributing to virulence. Proceedings of the National Academy of Sciences 116(16):8054–8059. https://doi.org/10.1073/pnas.1819481116

Huson DH, Beier S, Flade I, Górska A, El-Hadidi M, Mitra S, Ruscheweyh HJ, Tappu R (2016) MEGAN Community Edition - Interactive Exploration and Analysis of Large-Scale Microbiome Sequencing Data. PLOS Computational Biology 12(6):e1004957. https://doi.org/10.1371/journal.pcbi.1004957

Jones P, Binns D, Chang HY, Fraser M, Li W, McAnulla C, McWilliam H, Maslen J, Mitchell A, Nuka G, et al (2014) InterProScan 5: genome-scale protein function classification. Bioinformatics 30(9):1236–1240. https://doi.org/10.1093/bioinformatics/btu031

Li H (2013) Aligning sequence reads, clone sequences and assembly contigs with BWA-MEM. https://doi.org/10.48550/arXiv.1303.3997, URL http://arxiv.org/abs/1303.3997

Li H, Handsaker B, Wysoker A, Fennell T, Ruan J, Homer N, Marth G, Abecasis G, Durbin R, 1000 Genome Project Data Processing Subgroup (2009) The Sequence Alignment/Map format and SAMtools. Bioinformatics 25(16):2078–2079. https://doi.org/10.1093/bioinformatics/btp352

McKenna A, Hanna M, Banks E, Sivachenko A, Cibulskis K, Kernytsky A, Garimella K, Altshuler D, Gabriel S, Daly M, et al (2010) The Genome Analysis Toolkit: A MapReduce framework for analyzing next-generation DNA sequencing data. Genome Res 20(9):1297–1303. https://doi.org/10.1101/gr.107524.110

Minio A, Cochetel N, Vondras AM, Massonnet M, Cantu D (2022) Assembly of complete diploid-phased chromosomes from draft genome sequences. G3 Genes\textbackslashtextbarGenomes\textbackslashtextbarGenetics 12(8):jkac143. https://doi.org/10.1093/g3journal/jkac143

O'Leary NA, Wright MW, Brister JR, Ciufo S, Haddad D, McVeigh R, Rajput B, Robbertse B, Smith-White B, Ako-Adjei D, et al (2016) Reference sequence (RefSeq) database at NCBI: current status, taxonomic expansion, and functional annotation. Nucleic Acids Research 44(D1):D733–D745. https://doi.org/10.1093/nar/gkv1189

Paineau M, Mazet ID, Wiedemann-Merdinoglu S, Fabre F, Delmotte F (2022) The Characterization of Pathotypes in Grapevine Downy Mildew Provides Insights into the Breakdown of Rpv3, Rpv10, and Rpv12 Factors in Grapevines. Phytopathology® 112(11):2329–2340. https://doi.org/10.1094/PHYTO-11-21-0458-R

Paineau M, Minio A, Mestre P, Fabre F, Mazet ID, Couture C, Legeai F, Dumartinet T, Cantu D, Delmotte F (2024) Multiple deletions of candidate effector genes lead to the breakdown of partial grapevine resistance to downy mildew. New Phytologist 243(4):1490–1505. https://doi.org/10.1111/nph.19861

Quinlan AR, Hall IM (2010) BEDTools: a flexible suite of utilities for comparing genomic features. Bioinformatics 26(6):841–842. https://doi.org/10.1093/bioinformatics/btq033

Shumate A, Salzberg SL (2021) Liftoff: accurate mapping of gene annotations. Bioinformatics 37(12):1639–1643. https://doi.org/10.1093/bioinformatics/btaa1016

Simão FA, Waterhouse RM, Ioannidis P, Kriventseva EV, Zdobnov EM (2015) BUSCO: assessing genome assembly and annotation completeness with single-copy orthologs. Bioinformatics 31(19):3210–3212. https://doi.org/10.1093/bioinformatics/btv351

Tjeldnes H, Labun K, Torres Cleuren Y, Chyżyńska K, Świrski M, Valen E (2021) ORFik: a comprehensive R toolkit for the analysis of translation. BMC Bioinformatics 22(1):336. https://doi.org/10.1186/s12859-021-04254-w

Wingerter C, Eisenmann B, Weber P, Dry I, Bogs J (2021) Grapevine Rpv3-, Rpv10- and Rpv12-mediated defense responses against Plasmopara viticola and the impact of their deployment on fungicide use in viticulture. BMC Plant Biol 21(1):470. https://doi.org/10.1186/s12870-021-03228-7
